# Recombination in a natural population of the bdelloid rotifer *Adineta vaga*

**DOI:** 10.1101/489393

**Authors:** Olga A. Vakhrusheva, Elena A. Mnatsakanova, Yan R. Galimov, Tatiana V. Neretina, Evgeny S. Gerasimov, Svetlana G. Ozerova, Arthur O. Zalevsky, Irina A. Yushenova, Irina R. Arkhipova, Aleksey A. Penin, Maria D. Logacheva, Georgii A. Bazykin, Alexey S. Kondrashov

## Abstract

Sexual reproduction which involves alternation of meiosis and syngamy is the ancestral condition of extant eukaryotes. Transitions to asexual reproduction were numerous, but most of the resulting eukaryotic lineages are rather short-lived. Still, there are several exceptions to this rule including darwinulid ostracods^1,2^ and timema stick insects^3^. The most striking of them is bdelloid rotifers^4^–^6^, microscopic freshwater invertebrates which underwent an extensive adaptive radiation after apparently losing meiosis over 10 Mya. Indeed, both the lack of males in numerous bdelloid species and the lack of proper homology between chromosomes^6^ rule out ordinary sex. However, this does not exclude the possibility of some other mode of interindividual genetic exchange and recombination in their populations^7^. Recent analyses based on a few loci suggested genetic exchanges in this group^8,9^, although this has been controversial^10^. Here, we compare complete genomes of 11 individuals from the wild population of the bdelloid rotifer *Adineta vaga,* and show that its genetic structure, which involves Hardy-Weinberg proportions of genotypes within loci and lack of linkage disequilibrium between distant loci, is incompatible with strictly clonal reproduction. Instead, it can emerge only under ongoing recombination between different individuals within this species, possibly through transformation. Such a genetic structure makes the population immune to negative long-term consequences of the loss of conventional meiosis^11^, although this does not necessarily imply that interindividual genetic exchanges in *A. vaga* are directly maintained by natural selection.

---

The mode of reproduction of bdelloid rotifers, the most prominent of the ancient asexual “evolutionary scandals”^12,13^, has long been a subject of debate. Currently, there is a consensus that, due to the absence of males and lack of homologous chromosomes^6^, they cannot have conventional meiotic sex. Still, it remains unclear if they regularly engage in interindividual genetic exchanges by other means^8^–^10,14^.

To elucidate the mode of reproduction of the bdelloids, we sequenced whole genomes of 11 wild-caught individuals of the bdelloid rotifer *Adineta vaga* (Fig. 1a). For this, we established 11 clonal lineages, L1-L11, each started from a single rotifer matching the morphological criteria of *A. vaga*. Each sample was sequenced on Illumina HiSeq to the coverage of ∼40-100X (Methods; Supplementary Table 1). Additionally, we sequenced one of the lineages, L1, on the MiSeq platform, which allowed us to generate a high-quality *de novo* genome assembly for this lineage (Methods; Supplementary Table 2; Extended Data Fig. 1).

**Fig. 1.**
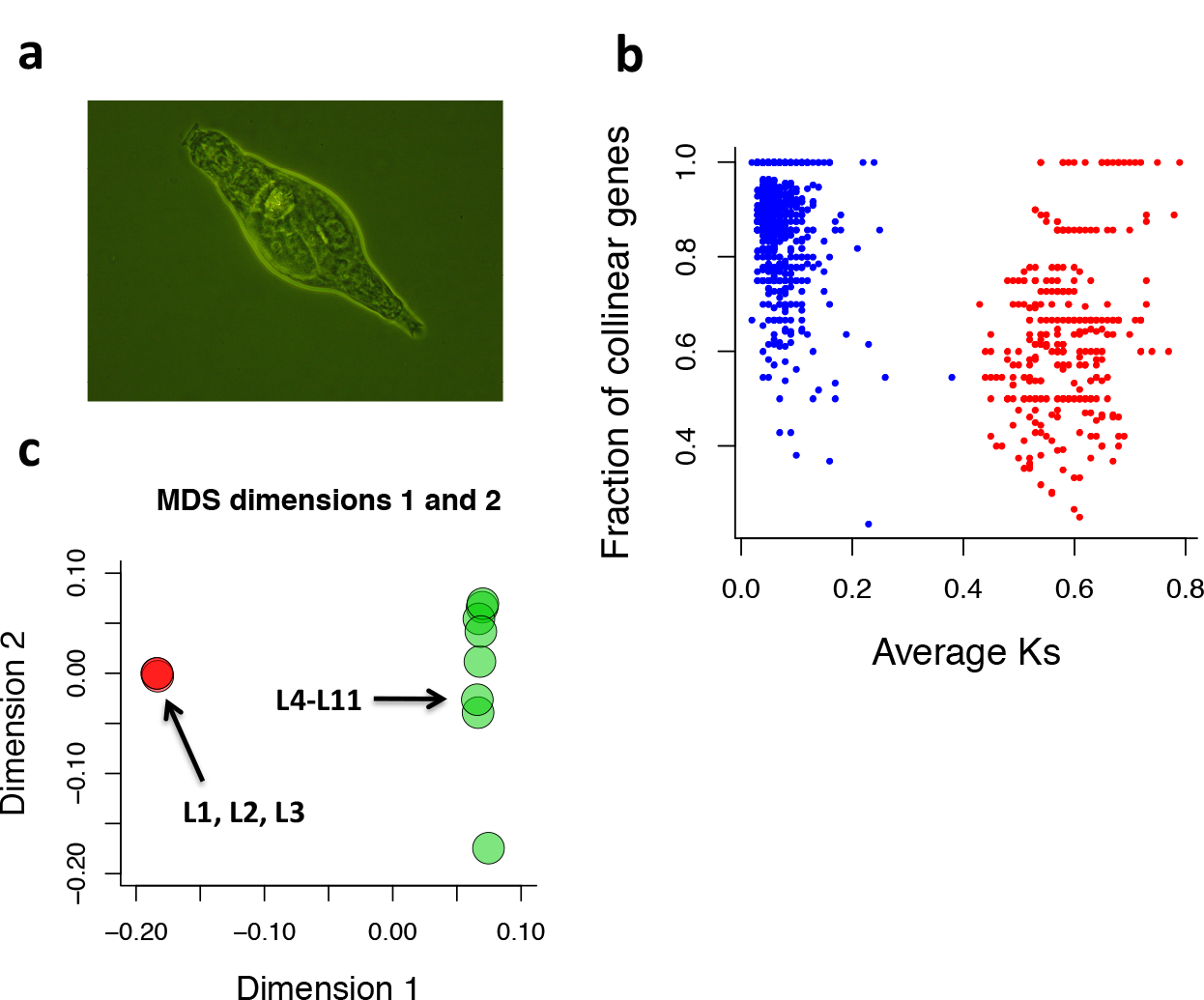
Whole-genome sequencing of 11 wild-caught individuals of the bdelloid rotifer *Adineta vaga*. **a,** Microphotograph of an *A. vaga* individual. From each of the 11 wild-caught rotifers matching the morphological description of *A. vaga*, a clonal lineage of over 1,000 individuals has been established and sequenced on the Illumina HiSeq to the coverage of ∼40-100X. **b,** Degenerate tetraploidy in the *A. vaga* L1 reference genome. The plot shows the mean number of synonymous substitutions between haplotypes per site (Ks) vs. the fraction of collinear genes for each syntenic block. Only blocks with 5 or more collinear genes are shown (n = 1,762). Blocks with average Ks values < 0.4 are in blue; those with average Ks ≥ 0.4 in red (see Supplementary Methods II). **c,** Genetic differentiation among the 11 sequenced individuals. The first two dimensions from multidimensional scaling analysis of pairwise genome-wide identities-by-state are shown. Genomes in red (L1-L3) and in green (L4-L11) were assigned to different clusters based on identity-by-state clustering analysis (P < 1e-100; see Supplementary Methods V).

The analysis of the obtained assembly, hereafter referred to as the reference genome, revealed the same patterns of genomic structure as those in the previously published genome of *A. vaga*^6^. Specifically, most of the genome could be assigned into pairs of collinear allelic regions representing two haplotypes, as well as into more distantly related clusters that probably arose from an ancient whole-genome duplication (Fig. 1b). We obtained a non-redundant haploid representation of the *A. vaga* genome (Methods; Supplementary Table 2; Supplementary Methods I) and mapped all the sequenced individual genomes against it (Methods; Supplementary Methods III).

Analysis of single-nucleotide differences between the sequenced individuals (Methods; Supplementary Table 4; Supplementary Methods IV) revealed presence of two genetic clusters (Fig. 1c; Supplementary Table 5; Supplementary Methods V). To minimize the potential effect of population structure, we focused the analysis on the 8 individuals (L4-L11) forming the larger cluster.

One of the key signatures of recombination in sexual populations is the decay of linkage disequilibrium (LD) with increasing physical distance between loci^15^. By contrast, in obligate asexuals, a complete LD is expected between different loci irrespective of the distance between them. We investigated the patterns of LD in the population of *A. vaga.* For each individual, we reconstructed the haplotypes by read-based phasing^16^ (Methods; Supplementary Tables 6, 7; Supplementary Methods VI). Contrary to what would be expected if the *A. vaga* population propagated strictly asexually, we found that LD within phased genome segments was much higher than that observed between segments (Fig. 2a; Methods; Supplementary Methods VII). Moreover, LD rapidly declined with the distance between polymorphic loci (Fig. 2a), reaching the level of r^2^ observed for different contigs at ∼2,800 nucleotides. Negative correlation of r^2^ with distance within individual segments^17^ as well as the sum of the distances^18^ and PHI tests^19^ also indicate that recombination is present (Fig. 3d; Methods; Supplementary Methods IX).

**Fig. 2.**
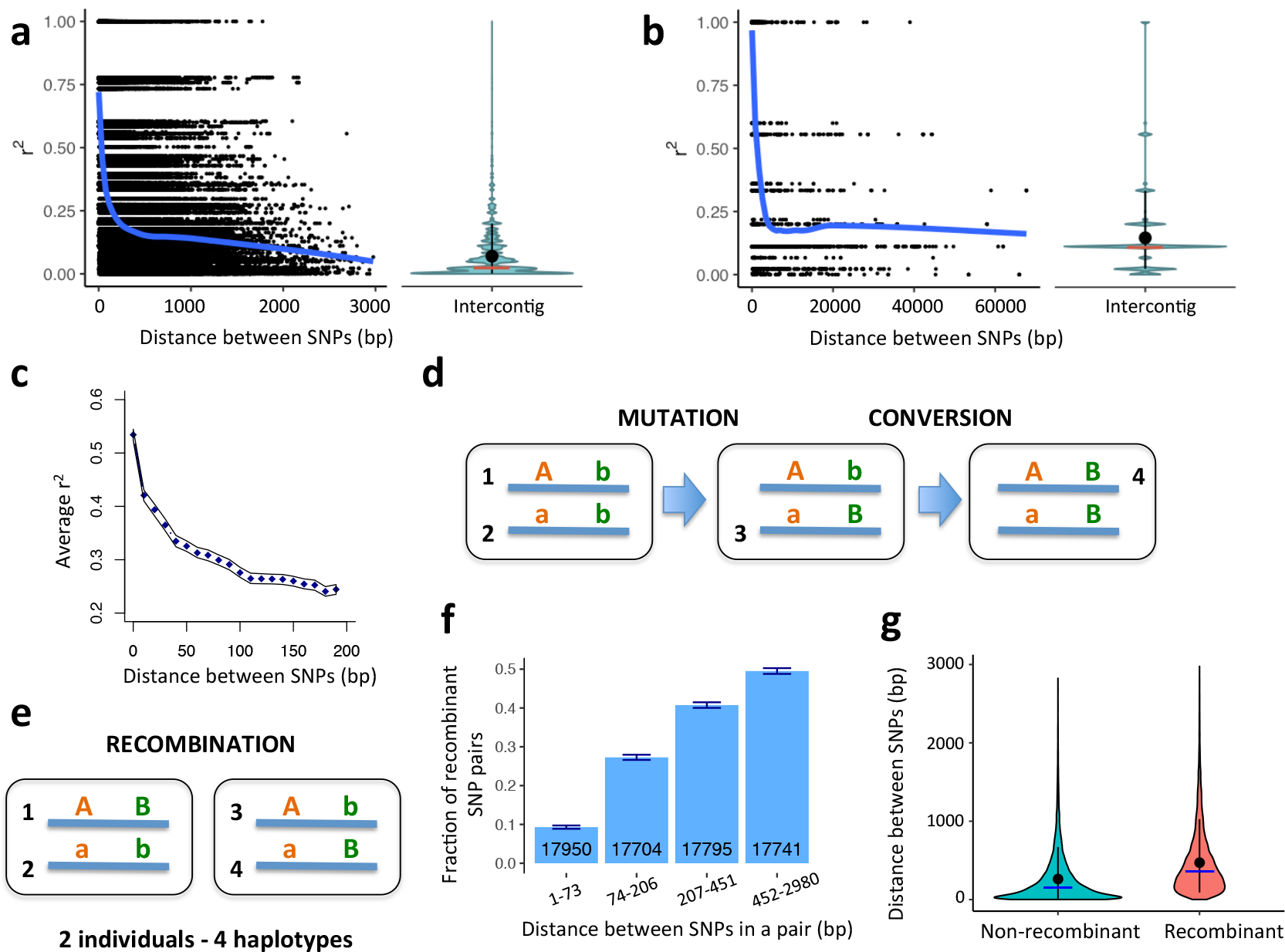
Decay of linkage disequilibrium with physical distance in *A. vaga*. **a, b,** Linkage disequilibrium is measured as r^2^. Decay of r^2^ with physical distance is estimated using phased haplotype data (**a**) and unphased genotype data (**b**). LOESS regression curves of r^2^ versus physical distance are shown in blue. Violin plots show the distributions of r^2^ values for the pairs of SNPs located on different contigs. Ends of the whiskers represent the 90% data range, with the mean and median values shown as a black dot and a red horizontal bar respectively. **a,** r^2^ was calculated using biallelic sites residing within the segments of the reference genome where haplotypes had been reconstructed for all the individuals forming the large cluster (L4-L11). **b,** Estimates of r^2^ based on the unphased genotype data were obtained using biallelic sites homozygous in all genomes forming the large cluster. **c,** Linkage disequilibrium is expressed as the squared correlation coefficient between genotypes. Squared correlation coefficients were computed for comparisons of 10,000 randomly drawn biallelic sites versus the remaining segregating biallelic sites. SNP pairs were subdivided by distance into bins of 10 bp; the displayed data are for the bins with SNP pairs separated by ≤ 200 bp. The confidence band is the 95% bootstrap confidence interval. **d,** Emergence of four haplotypes in the absence of genetic exchange and reciprocal recombination due to mutation and conversion. Boxes represent individuals. **e,** Schematic representation of a recombinant pair of positions which could not be the result of gene conversion. For a pair of SNPs, all four haplotypes are present in two individuals. Such pairs are regarded as passing the modified four-gamete test. **f,** Dependence of the fraction of site pairs passing the modified four-gamete test on the physical distance between the sites in a pair. For each pairwise combination of individuals, only those pairs of polymorphic sites that are simultaneously heterozygous in both individuals are considered. Error bars indicate exact 95% binomial confidence intervals. Site pairs meeting the requirements of the modified four-gamete test were subdivided into 4 distance bins with approximately equal numbers of cases. The total number of analyzed site pairs for each distance bin is shown at the base of the bars. Comparisons between all bin pairs were significant (P < 2e-16, Pearson Chi-square test, Holm correction for multiple comparisons). **g,** Violin plot showing the distribution of distances between sites in a pair for non-recombinant (n = 48,657) and recombinant (n = 22,533) pairs of sites. Whiskers indicate the 90% data range; mean and median distances for each group are shown with a black dot and a blue horizontal bar. Mean = 261.7 and median = 146 base pairs, non-recombinant; 470.6 and 353, recombinant. Mann-Whitney U test, P < 2.2e-16.

**Fig. 3.**
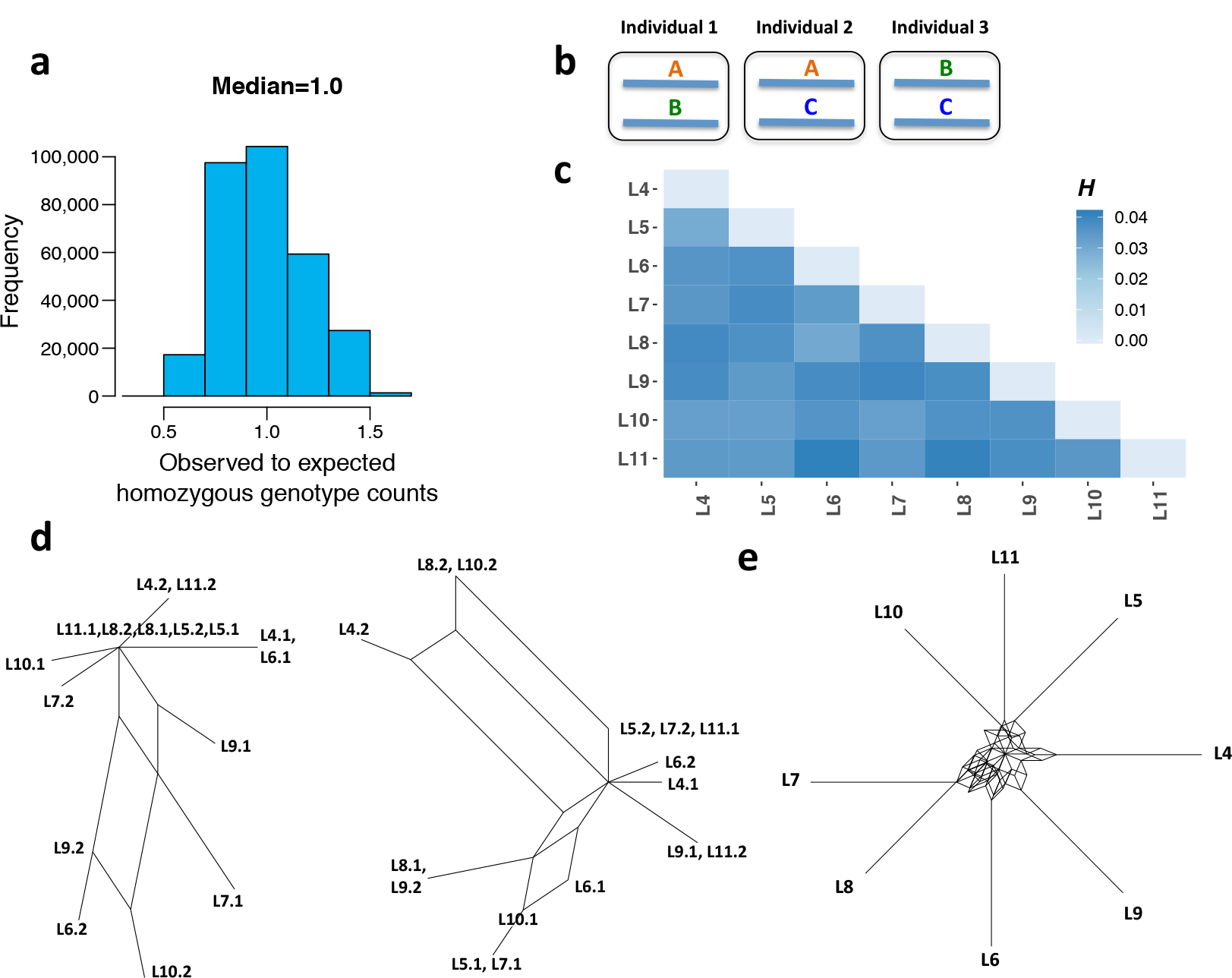
Population genomic analysis reveals signatures of genetic exchange in *A. vaga*. **a,** Distribution of observed-to-expected ratios for the numbers of genotypes homozygous for the major variant (individuals L4-L11, biallelic sites only, minor allele count ≥ 4, n = 307,049). **b,** A schematic representation of a triallelic site with segregating alleles A, B and C and all three their possible heterozygous combinations present among the sequenced individuals. **c,** Heatmap showing distribution of *H*-scores for pairs of *A. vaga* individuals. For each pair, the *H-*score reflects the probability to observe a site harboring all three heterozygotes as a result of hypothetical exchange between these two individuals (see Supplementary Methods XIII). **d,** Examples of split decomposition networks for two phased segments showing significant evidence for recombination according to the PHI test. Overall, the evidence for recombination was significant for 167 out of 262 segments (see Supplementary Methods IX). Indices 1 and 2 designate the two haplotypes of a single individual. **e,** Consensus network constructed from the neighbor-joining trees built using different allelic regions (n = 796) for the samples L4-L11. The network displays splits occurring at least in 10% of the individual trees with the length of the edges reflecting frequency of the splits (see Supplementary Methods X).

Consistent with the presence of recombination, the phylogenies of different genomic regions are incongruent. Indeed, the consensus network^20,21^ built from trees obtained for multiple regions has a reticulate shape suggestive of recombination (Fig. 3e; Extended Data Fig. 3c, d; Supplementary Methods X; Supplementary Note).

Even in the absence of recombination, the observed LD decay could arise artefactually. First, it might be caused by erroneous mapping of reads to paralogous regions. However, the decay persists in subsets of polymorphic loci covered by long blocks of collinear genes^22^ (Extended Data Fig. 2a), making this explanation unlikely. Second, it could arise due to phasing errors, including those arising from PCR template switching. To minimize the likelihood of this, we filtered the phased haplotypes aggressively (Methods; Supplementary Methods VI), and found that the rate of LD decay was virtually independent of the severity of filtering (Extended Data Fig. 2b). Still, residual phasing errors are conceivable. To ensure that the LD decay is not explained by them, we independently assessed it from unphased genotype data using two different approaches: inferring haplotypes from variable homozygous sites (Fig. 2b) or estimating LD directly from correlations between genotypes (Fig. 2c; Supplementary Methods VII). LD decay was observed in both these analyses. As these approaches do not rely on phasing, the observed decline of LD with distance between sites is not due to phasing errors.

Finally, LD decay could arise due to gene conversion between allelic regions. Gene conversion has been previously proposed to inflate the rate of LD decay between tightly linked loci in humans^23,24^, and it has been suggested to act within diploid loci of a single *A. vaga* individual^6^. In the classical Hudson’s four-gamete test^25^, the presence of all four possible types of haplotypes for a pair of biallelic polymorphic loci within a population is interpreted as evidence for recombination, because recurrent mutations are unlikely. However, a mutation followed by gene conversion would suffice to explain the presence of all four haplotypes without assuming genetic exchange between homologous regions (Fig. 2d). Nevertheless, the action of within-locus gene conversion can only produce a homozygous genotype from a heterozygous one, but not *vice versa*. Therefore, it cannot produce a pair of individuals, each heterozygous at two loci, carrying all four haplotypes (Fig. 2e; Supplementary Methods VIII); while such a pair can obviously arise through recombination involving genetic exchanges between individuals. We find that among the pairs of SNPs each heterozygous in two individuals, the fraction of those giving rise to all four possible haplotypes in these individuals is low when these SNPs are positioned nearby, but increases rapidly with the physical distance between SNPs (Fig. 2f, g; Supplementary Methods VIII). This observation is incompatible with the action of gene conversion as the sole cause of the LD decay (Supplementary Note).

Thus, we have to accept that there is some kind of genetic exchange between homologous regions. Still, conceivably, this exchange could involve only pairs of haplotypes residing within a single individual, without any transfer of genetic material between individuals.

To address this possibility, we analyzed the patterns of variation within individual biallelic SNPs. Under obligate asexual reproduction, two alleles at a locus accumulate mutations independently of each other. In a finite population, this creates a major excess of heterozygotes relative to the Hardy-Weinberg expectation^26,27^. By contrast, within-locus gene conversion obviously creates an excess of homozygotes. In our data, 97% of biallelic SNPs did not display significant deviations from the Hardy-Weinberg equilibrium (exact test, P > 0.05; Methods; Supplementary Methods XI), and this fraction was even higher when only the genomic regions with high-confidence ploidy were considered (97.4% and 97.5% for allelic regions and alleles respectively; Supplementary Table 8). The distribution of the observed-to-expected ratios for the number of homozygous genotypes was concentrated around 1 (median = 1.0, mean = 0.989; Fig. 3a; Supplementary Table 8; Supplementary Methods XI; Extended Data Fig. 3a, b), suggesting that the population is in Hardy-Weinberg equilibrium. In a strictly clonal population, this would imply that, for a particular set of parameters characterizing mutation and random drift, gene conversion occurs at a precisely specified rate. Because such fine-tuning of unrelated processes is unlikely, the observed Hardy-Weinberg proportions of diploid genotypes argue against obligate asexuality and suggest that genetic exchanges occur in *A. vaga*.

Independent evidence for genetic exchange between individuals is provided by triallelic SNPs^28^ carrying three out of four possible nucleotides. Triallelic SNPs can arise in sexual as well as in asexual organisms through multiple mutations affecting the same genomic site. At a polymorphic site with three alleles A, B and C, three heterozygous genotypes, A/B, A/C and B/C, are possible (Fig. 3b). Genetic exchanges between individuals will lead to frequent coexistence of all these genotypes. By contrast, in a population of asexuals, all three heterozygous genotypes can only arise through recurrent or back mutations, which are comparatively rare.

We identified 1,839 triallelic SNPs harboring all three possible heterozygotes among individuals L4-L11, compared to only 83.5 such sites expected due to recurrent or back mutations (P < 2.2e-16, one-sample Z-test for proportions; Supplementary Table 9; Supplementary Methods XII), a 22-fold enrichment. A similar excess was observed in the genomic regions with high-confidence ploidy, and was evident irrespective of whether only the individuals from the large cluster (L4-L11) or all sequenced individuals (L1-L11) were analyzed (Supplementary Table 9).

Rare heterozygous genotypes that likely arose from genetic exchanges are distributed almost uniformly among the individuals L4-L11, with different individuals possessing very similar numbers of such genotypes (Supplementary Table 10; Supplementary Methods XIII). To study in more detail how the genetic material has been exchanged in the *A. vaga* population, we designed a score *H* reflecting, for a pair of individuals, how likely it is that a site with all three heterozygous genotypes has emerged as a result of hypothetical exchange between these individuals (Supplementary Methods XIII). The values of *H* for different pairs L4-L11 were very similar (Fig. 3c; Supplementary Table 11), indicating that recombination signal is not driven by any particular pair of individuals. Together, these results argue against contamination as a source of the sites harboring three heterozygotes (Supplementary Methods XIII). More generally, they suggest a panmictic nature of the *A. vaga* population.

Overall, our data demonstrate that the genetic variation in the wild population of the bdelloid rotifer *A. vaga* was shaped by genetic exchanges between individuals and recombination, and is inconsistent with obligate asexuality. Given the evidence of horizontal gene transfer from non-metazoan species^29^–^31^ and indications of uptake of foreign DNA in bdelloids^32^, it is possible that intra-species genetic exchanges also occur through transformation^9^. An alternative possibility is a highly atypical version of meiosis observed in a few plants that involves segregation without crossing over (*Oenothera*-like meiosis) as hypothesized by Signorovitch *et al*. (ref. 8). To determine the mode of genetic exchange in *A. vaga,* we compared phylogenies of the two haplotypes harbored by each individual at different genomic loci. Transformation as well as *Oenothera*-like meiosis is expected to result in incongruent phylogenies of the two haplotypes^8^. However, unlike *Oenothera*-like meiosis, transformation should create different patterns of incongruence between haplotypes at different loci^8^. We observed multiple cases when the two haplotypes of a single individual clustered with haplotypes from different individuals (Extended Data Table 1; Fig. 4a, b, c, d; Methods). Such cases were detected in all individuals L4-L11 and supported different patterns of clustering of the two haplotypes of the same individual at different loci (Extended Data Table 1; Fig. 4a, b, c, d; Supplementary Table 12). This observation is consistent with the transformation scenario but is unlikely to result from *Oenothera*- like meiosis alone or from gene conversion (Supplementary Note). Another line of evidence against atypical meiosis as the main mode of genetic exchanges in *A. vaga* comes from highly correlated distances separating haplotypic counterparts for different pairs of individuals (Extended Data Fig. 6a, b, c; Supplementary Tables 13-15; Supplementary Methods XVII; Supplementary Note).

**Fig. 4.**
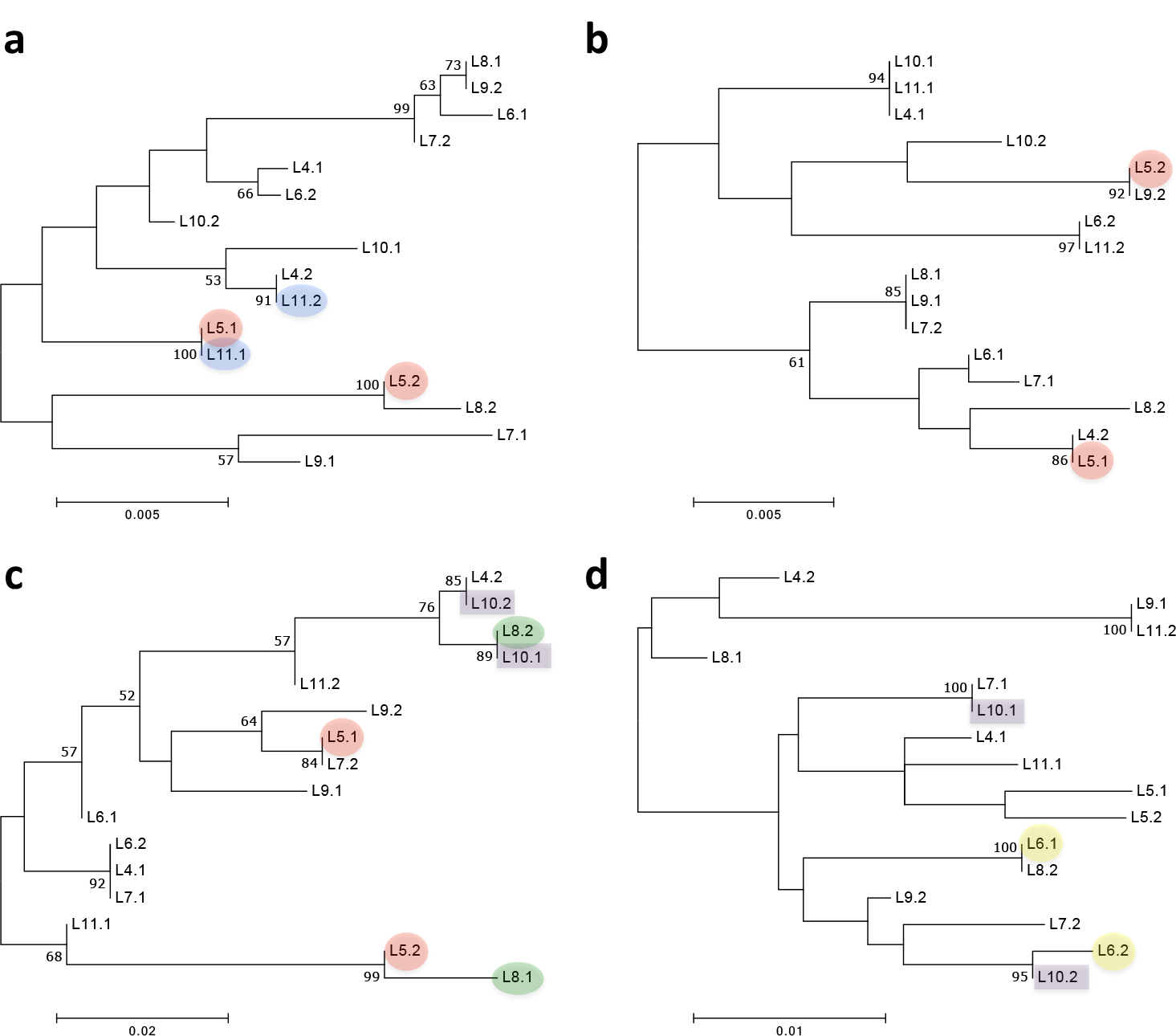
Phylogenetic analysis of haplotypes reveals signatures of interindividual genetic exchanges in *A. vaga*. **a, b, c, d,** Phylogenetic trees constructed for four phased segments exhibiting incongruent groupings of individuals with respect to two haplotypes. Overall, in 56 out of 262 segments, two haplotypes of at least one individual had their reciprocal closest counterparts in two different individuals (see Methods and Extended Data Table 1). The displayed phylogenies were constructed for four phased segments located on different contigs in the initial assembly and spanning 1,448 (**a**), 604 (**b**), 358 (**c**) and 1,198 (**d**) base pairs. Trees were built using maximum likelihood method in PhyML^41^ under the GTR+G model with 500 bootstrap trials. The bootstrap support values ≥ 50 % are shown next to branches. Only non-singleton SNPs that satisfied all filtering criteria and were simultaneously phased in L4-L11 were used to reconstruct phylogenies (n = 48, 21, 31 and 57 in **a, b**, **c** and **d** respectively); the remaining sites were treated as monomorphic (see Methods). Indices 1 and 2 designate the two haplotypes of a single individual. In these four segments, the two haplotypes of the same individual clustered with haplotypes from two different individuals with high bootstrap support values (>80%) for L5 (**a, b, c**; red), L6 (**d**; yellow), L8 (**c**; green), L10 (**c, d**; violet) and L11 (**a**; blue).

Under transformation scenario, the rate of LD decay would depend on the transformation rate. The probability of recombination between adjacent sites caused by homologous replacement of DNA segments would be determined by the rate of transformation, independently of the length of the transferred segments (Supplementary Methods XIV). Based on the values of recombination fractions^33,34^ assessed from the data, *A. vaga* might experience on average ∼1 transformation event per generation (Supplementary Methods XIV, XV). The LOESS estimates of r^2^ reach the values close to the intercontig level already at 1,500-2,000 nucleotides (Fig. 2a), suggesting that the characteristic size of transferred segments should also be at least 1,500-2,000 nucleotides (Supplementary Methods XIV).

Recombination is more frequent in GC-poor regions of the *A. vaga* genome (Extended Data Fig. 4a, b, c; Extended Data Fig. 5a, b; Methods; Supplementary Methods XVI). A similar pattern has been observed in *Arabidopsis*^35^, although a positive relationship between recombination rate and GC-content, attributed to gene conversion, appears to be more common^36,37^. Perhaps, fragile GC-poor regions are more prone to incorporation of external DNA in *A. vaga*. Further, similarly to what has been observed in other species^38^, exons in *A. vaga* recombine more frequently than introns (Extended Data Fig. 4d; Supplementary Methods XVI).

In summary, despite near-certain lack of conventional meiosis and syngamy, the natural population of *A. vaga* is essentially amphimictic, lacking both within- and between-locus associations between alleles. Panmixis by transformation has been previously reported in bacteria^7^, but was unheard of in eukaryotes. Although the nature of evolutionary advantage of sex remains unclear^39^, recombination in *A. vaga*, irrespective of its mechanism, is likely to protect it from the long-term costs of abandoning sex such as Muller’s ratchet. However, this does not necessarily imply that recombination is favored by natural selection due to its genetic effects^40^.

## Supporting information

**Extended Data Fig. 1.**
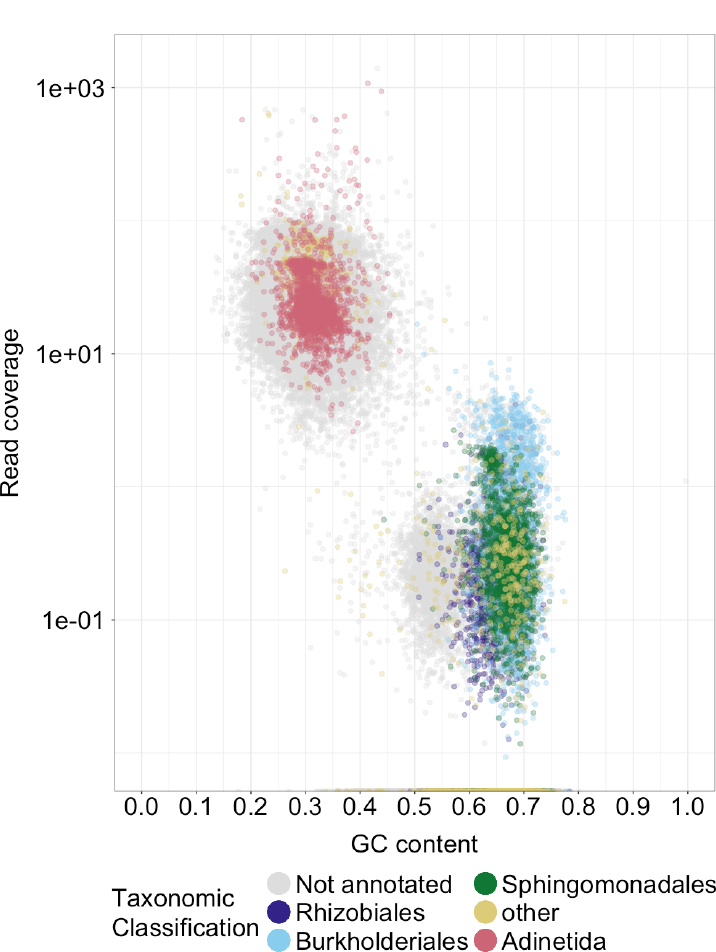
Taxonomic classification of contigs from the initial assembly of the *A. vaga* genome. GC-content versus read coverage for the contigs from the initial assembly of lineage L1 from MiSeq reads. Read coverage was determined from the HiSeq reads which were obtained from a separate library and not used for assembly (see Methods). Contigs are colour coded based on taxonomic classification^42^ of their best BLAST hit to nt database with E-value cutoff set to 1e-5. Only taxonomic annotations ascribed to at least 6% of the contigs are displayed; unannotated contigs and contigs representing less abundant annotations are shown in grey and yellow respectively.

**Extended Data Fig. 2.**
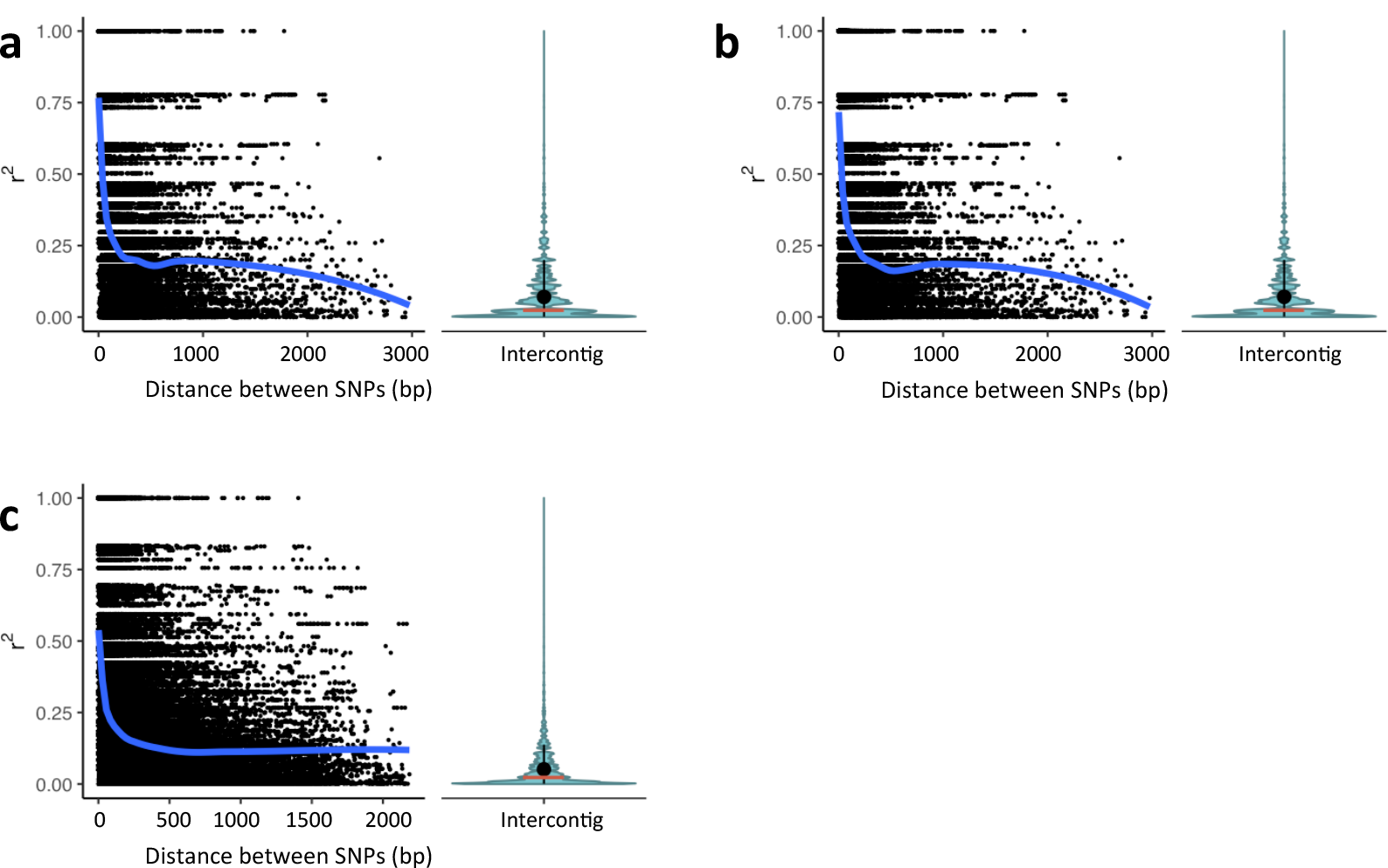
Decay of linkage disequilibrium with physical distance among the sequenced *A. vaga* individuals is robust to erroneous read alignment, phasing errors and effects of population structure. **a, b, c,** Linkage disequilibrium is measured as r^2^. Decay of r^2^ with physical distance is estimated using phased haplotype data. LOESS regression curves of r^2^ versus physical distance are shown in blue. Violin plots show the distributions of r^2^ values for the pairs of SNPs located on different contigs. Ends of the whiskers represent the 90% data range, with the mean and median values shown as a black dot and a red horizontal bar respectively. **a, b,** Decay of linkage disequilibrium with physical distance among the individuals L4-L11. r^2^ was estimated using biallelic sites residing within the segments of the *A. vaga* genome where haplotypes had been reconstructed for all the individuals forming the large cluster (L4-L11). **a,** r^2^ was calculated using biallelic sites from the phased dataset 1 residing within the allelic regions of the *A. vaga* genome (see Supplementary Methods II). **b,** r^2^ was calculated using biallelic sites from the stringently filtered phased dataset 2 (see Supplementary Methods VI). **c,** Decay of linkage disequilibrium with physical distance among the individuals L1-L11. r^2^ was calculated using biallelic sites from the phased dataset 1 residing within the segments of the *A. vaga* genome where haplotypes had been reconstructed for the complete set of individuals L1-L11.

**Extended Data Fig. 3.**
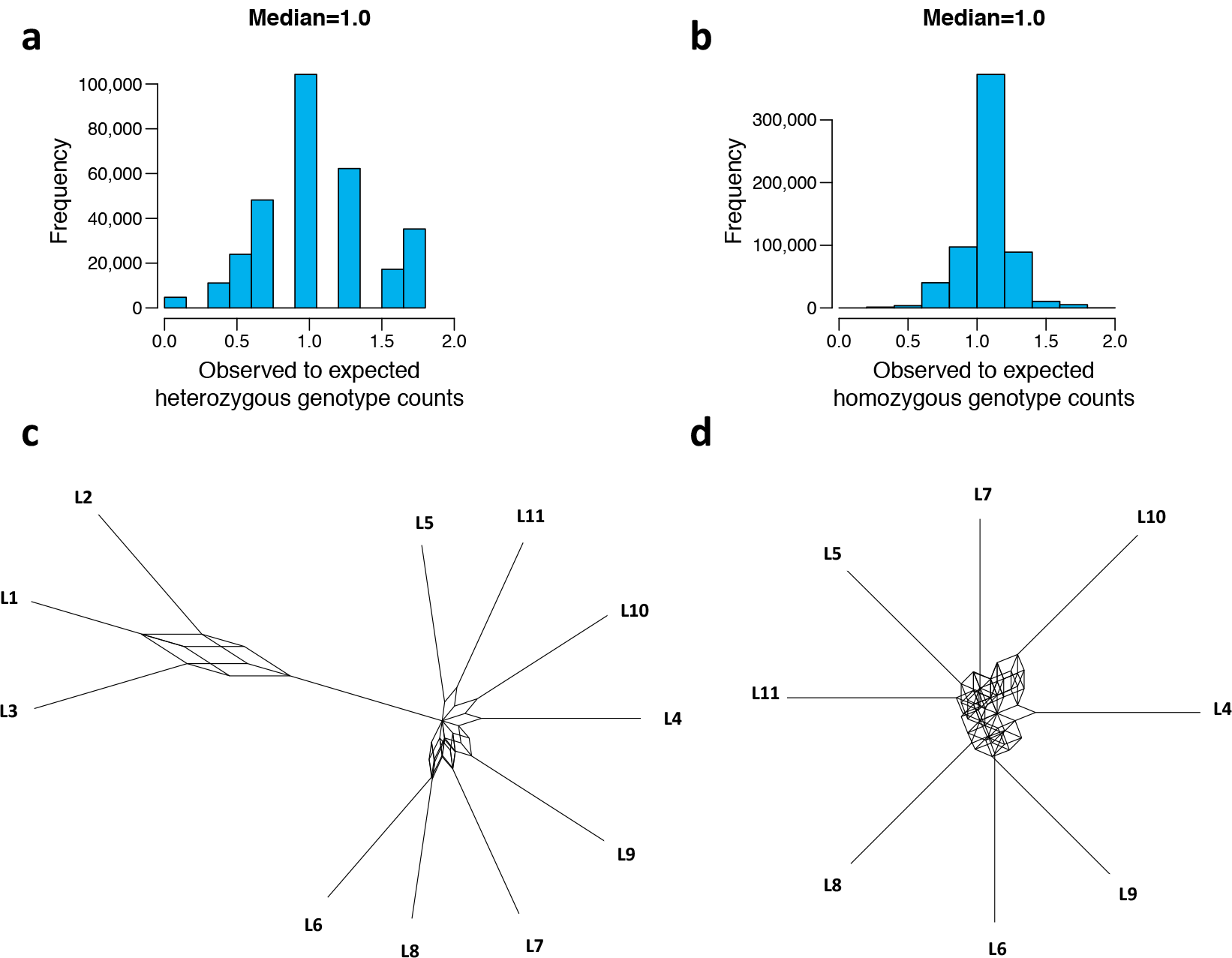
Population genomic analysis reveals signatures of genetic exchange in *A. vaga*. **a,** Distribution of observed-to-expected ratios for the numbers of heterozygous genotypes (individuals L4-L11, biallelic sites only, minor allele count ≥ 4, n = 307,049). **b,** Distribution of observed-to-expected ratios for the numbers of genotypes homozygous for the major variant (individuals L1-L11, biallelic sites only, minor allele count ≥ 4, n = 620,102). **c, d,** Consensus networks constructed from the neighbor-joining trees built using different allelic regions (**c**, n = 832) or allelic genes (**d**, n = 680) of the *A. vaga* genome for individuals L1-L11 (**c**) or L4-L11 (**d**). The networks display splits occurring at least in 10% of the individual trees, with the length of the edges reflecting frequency of the splits (see Supplementary Methods X).

**Extended Data Fig. 4.**
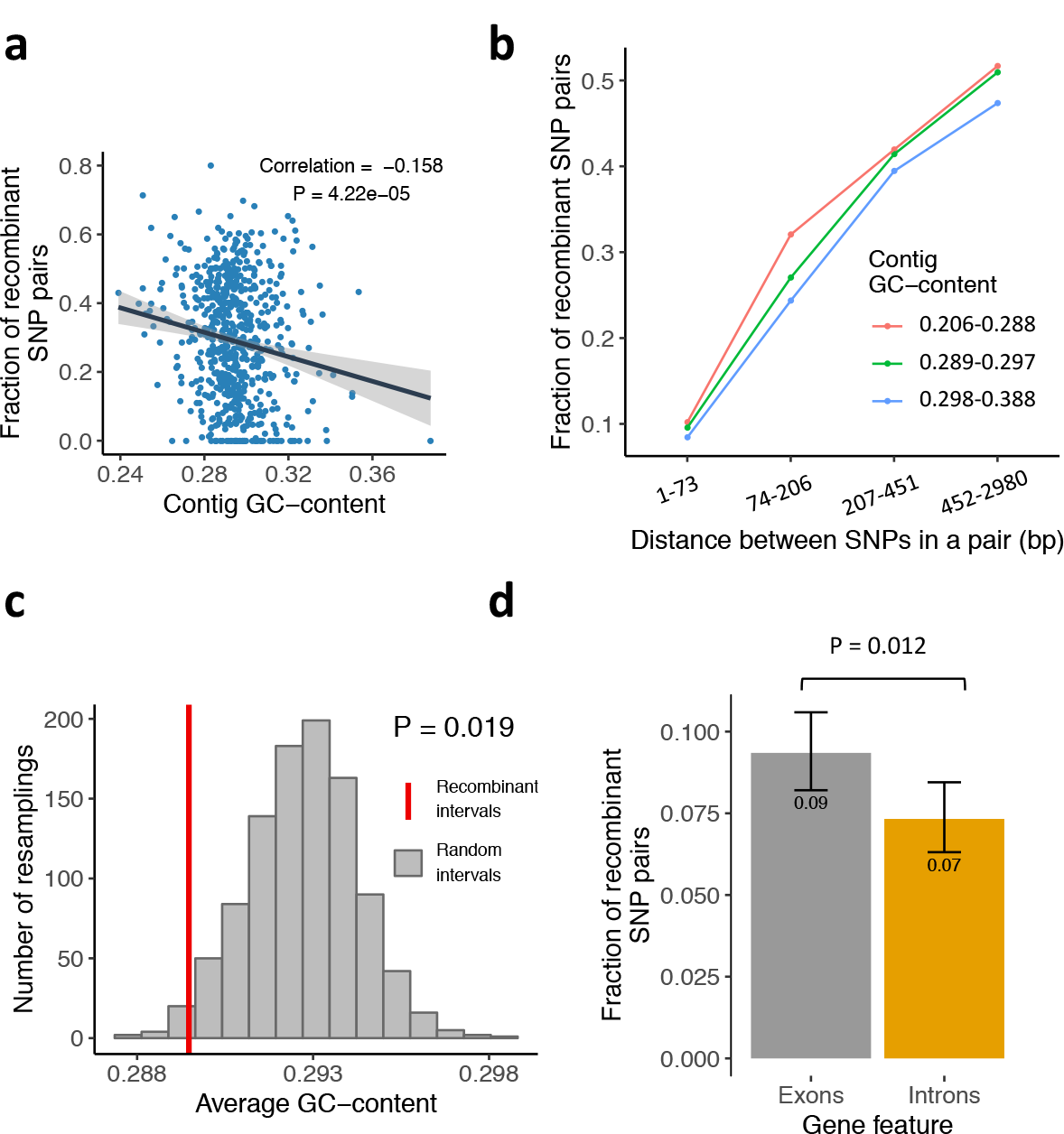
Features of recombination process in *A. vaga*. a, b, c, Recombination is targeted towards GC-poor regions of the *A. vaga* genome. **a,** Negative correlation between the contig GC-content and the fraction of recombinant SNP pairs. The line and the shaded area represent the best fit and the 95% confidence band respectively. **b,** Fraction of SNP pairs passing the modified four-gamete test as a function of distance between SNPs in a pair and contig GC-content. The data are the same as in Fig. 2f, g. For the 95% binomial confidence intervals and P values, see Extended Data Fig. 5b. **c,** GC-content of genomic intervals residing between recombinant pairs of sites compared to GC-content of random genomic intervals. The distribution was obtained by sampling 1,000 times random genomic intervals matched for number (n = 1,014), length and contig identity to the recombinant intervals. The one-sided P value was calculated as the fraction, among 1,000 sets of random intervals, of those with the average GC-content lower or equal to that of the recombinant intervals (see Supplementary Methods XVI). **d, Recombination is targeted towards exons in *A. vaga*.** Fraction of recombinant SNP pairs among the pairs of heterozygous SNPs residing within 200 bp of each other for exons and introns (pairs of SNPs in exons were matched for distribution of distances between the sites in a pair to SNPs in introns; n = 2,375 for both exons and introns; mean distance between SNPs in a pair = 33 bp for both exons and introns). Error bars indicate exact 95% binomial confidence intervals. The exact fractions are displayed below the error bars. P value is from two-sample Z-test for proportions (see Supplementary Methods XVI).

**Extended Data Fig. 5.**
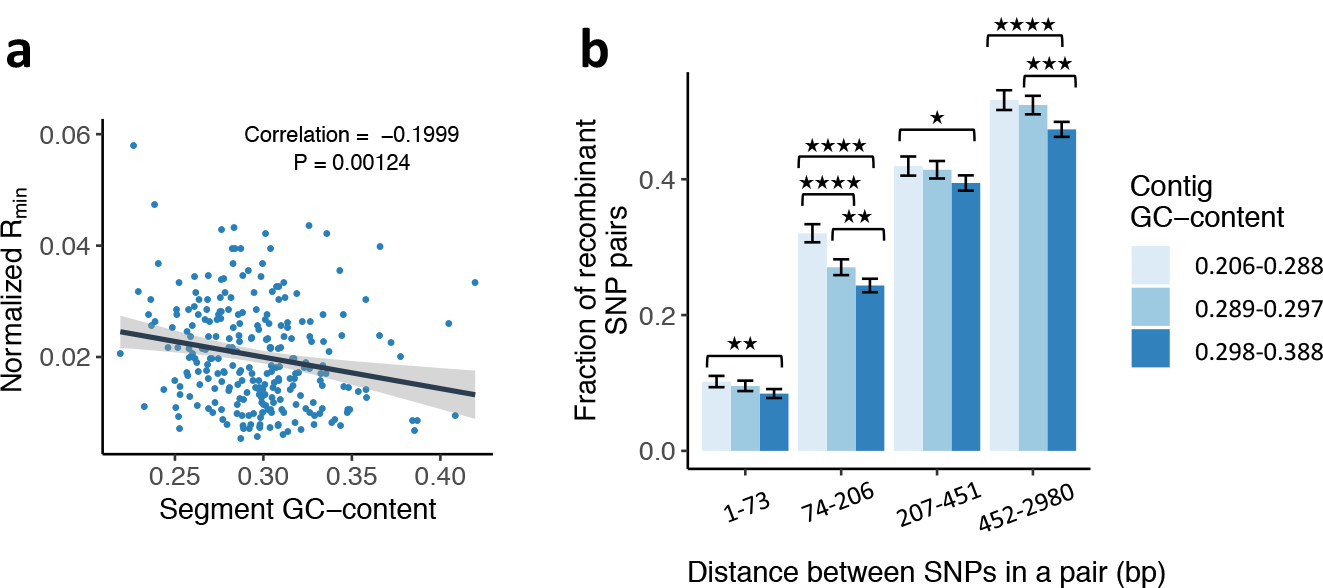
Recombination is targeted towards GC-poor regions of the *A. vaga* genome. **a,** Correlation plot showing the relationship between normalized minimum number of recombination events (R_min_) and GC-content of the genomic segment. The line and the shaded area represent the best fit and the 95% confidence band, respectively. Minimum numbers of recombination events were estimated for individual phased genomic segments harboring no less than 20 phased non-singleton SNPs in L4-L11 (n = 258; see Supplementary Methods XVI). **b,** Barplot showing dependence of the fraction of SNP pairs passing the modified four-gamete test on physical distance between SNPs in a pair and contig GC-content. The data are the same as in Extended Data Fig. 4b and Fig. 2f, g. Error bars indicate exact 95% binomial confidence intervals. Asterisks indicate significantly different fractions of recombinant SNP pairs for the comparisons of groups differing in GC-content within the same bin of inter-site distance. * P < 0.05, ** P < 0.01, *** P < 0.001, **** P < 0.0001. All pairwise comparisons between groups from different distance bins were significant at P < 0.0001. P values are from Pearson Chi-square test with Holm correction for multiple comparisons.

**Extended Data Fig. 6.**
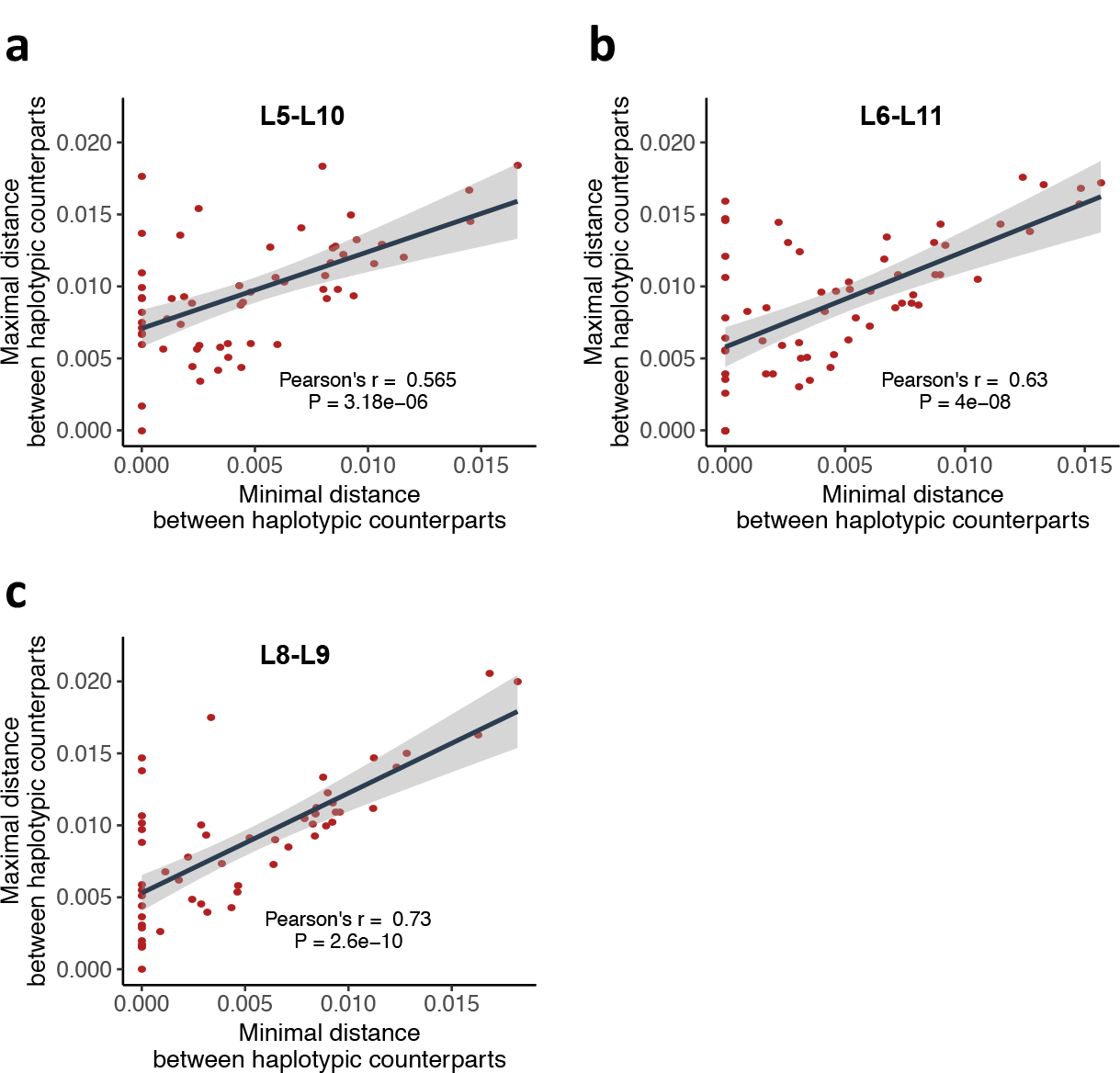
Correlation between minimal and maximal distance separating haplotypic counterparts for three pairs of individuals. **a, b, c,** Correlation plots showing the relationship between minimal and maximal distance separating haplotypic counterparts for pairs of individuals L5-L10 (**a**), L6-L11 (**b**) and L8-L9 (**c**). Points represent different phased segments. Distances are expressed as proportions of nucleotide differences. Pearson’s r between minimal and maximal distance separating haplotypic counterparts in different phased segments and the corresponding P values are shown. The line and the shaded area represent the best fit and the 95% confidence band, respectively. Distances were computed for the segments of the *A. vaga* genome simultaneously phased in all individuals L4-L11 and carrying at least 20 non-singleton SNPs. For each comparison of individuals, we utilized only those segments where 4 haplotypes from two individuals could be unambiguously assigned into 2 pairs, each pair comprising a haplotype from the first individual and its counterpart from the second individual. Haplotypes assigned to the same pair are likely to share a recent common ancestor. Only those variable positions that passed all filtering steps and were simultaneously phased in L4-L11 were taken into account to compute distances (see Supplementary Methods XVII). For correlations between all pairs of individuals L4-L11 see Supplementary Table 13.

**Extended Data Table 1.** Incongruent groupings of the two haplotypes in individuals L4-L11. For each individual, we computed the number of phased segments such that the reciprocal closest counterparts of the two haplotypes were found in two different individuals (see Methods). Only incongruent groupings of the two haplotypes with strong bootstrap support (≥70%) were considered. The numbers of identified incongruent segments along with the numbers of different patterns of incongruence observed for each individual, L4-L11, are shown. For each individual, each unique pair of other individuals harboring reciprocal closest counterparts of its two haplotypes at least at one locus constitutes a separate pattern of incongruence. The patterns of incongruence observed for each individual are listed, with the number of segments for which each pattern was observed given in parentheses. For this analysis, we used only the segments of the *A. vaga* genome simultaneously phased in all individuals L4-L11 and carrying at least 20 non-singleton SNPs. In total, out of the 262 analyzed phased genomic segments, 56 exhibited incongruent groupings of the two haplotypes at least in one individual with strong bootstrap support. Overall numbers of different patterns of incongruence identified in analyzed segments are shown in Supplementary Table 12.

| Individual | Number of |  |  | Observed patterns of incongruence |
| --- | --- | --- | --- | --- |
|  | Analyzed phased segments | Incongruent phased segments | Different patterns of incongruence |  |
| **L4** | 262 | 6 | 4 | L5-L11 (1), L6-L11 (3), L7-L10 (1), L10-L11 (1) |
| **L5** | 262 | 8 | 7 | L4-L7 (1), L4-L9 (1), L4-L10 (2), L6-L11 (1), L7-L8 (1), L8-L11 (1), L9-L10 (1) |
| **L6** | 262 | 10 | 8 | L4-L10 (1), L5-L8 (2), L5-L9 (1), L5-L11 (1), L7-L10 (1), L8-L10 (1), L8-L11 (2), L9-L11 (1) |
| **L7** | 262 | 6 | 5 | L4-L8 (2), L4-L9 (1), L5-L9 (1), L9-L10 (1), L9-L11 (1) |
| **L8** | 262 | 13 | 8 | L4-L5 (1), L5-L7 (1), L5-L10 (2), L6-L9 (2), L6-L11 (4), L7-L11 (1), L9-L10 (1), L9-L11 (1) |
| **L9** | 262 | 11 | 8 | L4-L8 (1), L5-L7 (2), L5-L10 (1), L5-L11 (1), L6-L11 (1), L7-L8 (1), L7-L10 (1), L8-L11 (3) |
| **L10** | 262 | 10 | 8 | L4-L8 (1), L4-L9 (1), L4-L11 (2), L5-L6 (2), L5-L7 (1), L6-L7 (1), L7-L8 (1), L7-L11 (1) |
| **L11** | 262 | 8 | 6 | L4-L5 (1), L4-L7 (1), L4-L9 (2), L5-L8 (2), L6-L10 (1), L7-L9 (1) |

## Methods

### Establishment of clonal cultures of *A. vaga*

We collected rotifers from clumps of moss which grew on trunks of aspen *Populus tremula* at height 120-170 cm near Biostation Lake Glubokoe in Ruza district of the Moscow region, Russia, and in the vicinity of village Shilovo in the Manturovo district of the Kostroma region, Russia. Individual rotifers were isolated and identified to the species level on the basis of morphological criteria^43^. Clonal cultures were started from individuals identified as *Adineta vaga*, which were rinsed in Milli-Q water and transferred to 96-well cell-culture plates. To confirm that just a single individual was transferred to each well, plates were visually inspected daily for the next 3 days after inoculation. When a culture reached the size of ∼30 individuals, it was transferred to a separate Petri dish containing Milli-Q water. Cultures were kept at 15-20 °C and fed *E. coli* (DH5α strain) grown overnight at 37°C in LB medium. When the total number of rotifers in the dish reached ∼1,000, they were harvested for DNA extraction. In total, 11 cultures of *A. vaga*, L1-L11, were established. The species identity of cultures L1-L11 was additionally confirmed by the number of pharyngeal teeth in the rake organ (four U-shaped hooks on each side of the mouth), a distinctive feature of *A. vaga*.

### DNA extraction

Total DNA was isolated on Promega Wizard SV Genomic DNA Purification Kit columns (Promega, United States) according to the manufacturer’s protocol.

### Library preparation

DNA was fragmented using Covaris S2 sonicator. Libraries were prepared using TruSeq DNA sample preparation kit (Illumina) following the protocol, with the following modification: for ligation, we used adapters from the TruSeq RNA sample preparation kit, as they have lower concentration and are thus more suitable for low-input samples.

### Genomic DNA sequencing

We sequenced the genomic DNA of 11 *A. vaga* cultures to the coverage of 40-100X (see Supplementary Table 1) with Illumina paired-end libraries on the Illumina HiSeq platforms 2000 or 2500 using TruSeq SBS kits v.3 or v.4 respectively. The exact numbers of paired-end reads generated for each culture and the resulting coverage are presented in Supplementary Table 1.

Additionally, we sequenced one of the lineages, L1, on the MiSeq platform to produce a high-quality *de novo* genome assembly (see section «Reference genome assembly and filtration»). To minimize any bias in the way different samples were analyzed, the MiSeq reads obtained for L1 were used only for assembly of the *A. vaga* reference genome. Variant calls for the lineage L1 were produced in the same way as for the rest of the samples, using alignments of 2×100 bp HiSeq reads.

### Removal of adapter sequences and quality trimming

We used Trimmomatic^44^ V0.33 to remove adapter sequences and perform quality trimming on raw sequencing reads. The parameters for quality trimming were set to “LEADING:20 TRAILING:20 SLIDINGWINDOW:5:20”. Reads shorter than 50 bp were discarded. The numbers of reads left after adapter and quality filtering for each individual are presented in Supplementary Table 1. We performed quality control checks on the reads with FastQC v0.11.3 (https://www.bioinformatics.babraham.ac.uk/projects/fastqc/).

### Reference genome assembly and filtration

The first genome of a bdelloid rotifer, *A. vaga,* was published in 2013^6^. However, we found that the sequence identity of our *A. vaga* genomes to the published genome was only ∼88% (Supplementary Table 3), which prevented us from using the published genome as a reference. This finding is in line with previous reports showing that morphological species of bdelloid rotifers frequently comprise complexes of divergent cryptic species^45^.

To obtain a genome assembly which could be used as a reference, we additionally sequenced one of our cultures, L1, on the MiSeq platform, which made it possible to generate a high-quality *de novo* genome assembly (Supplementary Table 2). For this purpose, we generated a separate Illumina library for L1 and sequenced it on the MiSeq system with Miseq reagent kits v.2 or v.3. Three sequencing runs were performed, yielding the totals of 10,172,970 (2×301 bp), 15,397,651 (2×251 bp) and 20,061,190 (2×261 bp) reads. Trimming of low-quality bases and adapter removal was performed using Trimmomatic^44^ V0.33 with the parameters for quality trimming set to “LEADING:20 TRAILING:20 SLIDINGWINDOW:5:20”. Reads shorter than 50 bp were discarded. The total of 34,403,183 reads left after these steps were used to produce a *de novo* genome assembly.

Assembly of the L1 genome was carried out with SPAdes^46^ (version: 3.6.0) and based on the MiSeq reads. SPAdes was run in diploid mode (--diploid option) without preliminary read error correction (--only-assembler option). K-mer sizes were set to: -k 21,33,55,77,99,127.

The initial assembly had an N50 of 18 kilobases and contained 233.8 Mbp of sequence in 51,852 contigs longer than 500 bp (Supplementary Table 2).

A bimodal distribution of the GC-content in the contigs suggested presence of bacterial contamination. However, contaminant-derived contigs were easily distinguished from the target *A. vaga* contigs by a significantly lower coverage and higher GC-content (Extended Data Fig. 1). Taxonomic classification of the initial contigs was performed with the Blobology pipeline^42^ according to best BLAST^47^ hit of a contig to nt database with E-value cutoff set to 1e-5. Taxon annotated GC-coverage plots^42^ were created with the modified R script (makeblobplot.R) provided as a part of the Blobology pipeline. Coverage of contigs was estimated from alignments of the MiSeq reads used in the assembly as well as from alignments of the HiSeq reads obtained from sequencing of the initial Illumina library constructed for the lineage L1.

The resulting taxon annotated GC-coverage plots were used to partition the contigs into sets of target and contaminant contigs (Extended Data Fig. 1). Assessing coverage of the contigs from the HiSeq reads, which were not used in the assembly, resulted in better discrimination between the target and contaminant contigs (not shown).

To filter out contaminant sequences, we first removed from the assembly all contigs with GC-content ≥ 0.5 or coverage with HiSeq reads < 4. To ensure that the final assembly comprised only contigs truly originating from *A. vaga*, we performed a BLAST^47^ search with the pre-filtered set of contigs against the published *A. vaga* genome^6^. BLAST searches were performed with *blastn* from BLAST+ 2.2.31 with the following parameters: -evalue 1e-10 -outfmt “6 qseqid sseqid pident length mismatch gapopen qstart qend sstart send evalue bitscore qlen slen” -task dc-megablast.

Only contigs with at least one dc-megablast hit to the published assembly with E-value ≤ 1e-100 and the minimal alignment length of 500 base pairs were retained. The resulting filtered assembly had an N50 of 22 kilobases, with 19,068 contigs longer than 500 bp covering 197 megabases of sequence, comparable with 218 megabases reported for the published *A. vaga* assembly^6^. Summary assembly statistics for the initial and filtered sets of contigs are shown in Supplementary Table 2. The filtered assembly was used as a reference in all subsequent analyses. The assembly statistics were generated with QUAST^48^ v5.0.0.

### Construction of non-redundant haploid sub-assembly

Due to high heterozygosity, the two haplotypes of the *A. vaga* genome assemble into separate contigs at the majority of loci^6^. Still, in a substantial portion of the genome, the two haplotypes collapse into a single contig, leading to a mosaic organization of the assembly with alternating ploidy levels. To reduce redundancy of the assembly and to ensure that only truly diploid loci are analyzed, we obtained a reduced haploid representation of the *A. vaga* genome.

Briefly, we searched for the pairs of reciprocally highly similar genomic segments within the assembly, discarding genomic regions without haplotypic counterparts and non-reciprocal best matches. From each pair of reciprocal best matches we retained only a single segment. This was achieved by using *blastn*^47^ from BLAST+ 2.2.3, blastz^49^ and single_cov2 commands from the all_bz^50^ v.15 program and custom Perl code. See Supplementary Methods I for details. The resulting haploid sub-assembly spanned 76,679,421 base pairs (Supplementary Table 2), suggesting that at least ∼80% of the initial assembly is represented by two haplotypes.

### Annotation of protein-coding genes

We predicted protein-coding genes in the filtered diploid assembly of the *A. vaga* genome using AUGUSTUS^51^ and GeneMark.ES^52^. Intron and transcribed region hints for AUGUSTUS were prepared with STAR aligner^53^. For this purpose, RNA-seq reads from http://www.genoscope.cns.fr/adineta/data/ were mapped on the initial assembly with strict mapping parameters. The list of putative splice junctions (a total of 119,058 suggested intron boundaries) was obtained taking into account only uniquely mapped reads (16% of the available RNA-seq reads, 21 million reads). Coverage profile for all mapped reads (+23% of reads that mapped to multiple loci, 61 million of total mapped reads) was assessed with a custom script using alignments produced by STAR. To compile the final set of intron and exon part hints for AUGUSTUS, we combined predictions of splice junctions with the coverage profile and filtered poorly supported junctions using a custom Python script (splice junction support cut-off > 1 unique reads, exonpart support cut-off > 5X coverage). GeneMark.ES was run to obtain a set of initial gene predictions. These predictions were ranked by *blastp* alignment quality (E-value, query coverage, gaps, positives), and 500 top-scoring models were used to generate the training set for AUGUSTUS. Species model was trained and then used to generate the final hinted predictions with AUGUSTUS. All predicted models (from AUGUSTUS and from GeneMark.ES) were combined together and scored according to the *blastp* alignment score and support from RNA-seq reads. To get rid of chimeric genes and mispredicted gene fragments, best gene models were selected at each locus. The initial set of predictions comprised 78,303 gene models originating from 75,877 loci.

We performed a quality check on the initial gene predictions, discarding gene models with putative annotation errors and those likely corresponding to incomplete genes. First, we removed annotations at the contig boundaries, which left us with 72,406 out of 78,303 gene models. Next, we checked CDS regions of each gene model, excluding a gene model from further analyses if the length of the corresponding CDS was not in multiple of three (n = 6,919).

We further filtered out gene models if the corresponding CDS carried a premature stop codon (n = 691) or was lacking a canonical termination codon (n = 1,426). A total of 63,370 gene models originating from 61,531 loci (genes) remained after these filtering steps. For each locus, the longest transcript among those that passed all filters was retained, providing a total of 61,531 gene models which were used in downstream analyses.

We transferred the filtered gene models predicted in the diploid assembly to the coordinate system of the haploid sub-assembly, retaining only those gene models that were fully contained within the haploid segments. Gene models partially overlapping with haploid segments were discarded. This procedure yielded 23,802 gene models with coordinates mapped to the coordinate system of the haploid sub-assembly. The filtering and conversion of gene coordinates between assemblies were performed with custom Perl scripts.

### Identification of allelic regions and allelic genes

To obtain a subset of the *A. vaga* genome with high-confidence ploidy, we identified genomic regions that could be assigned into pairs of highly similar segments with conserved gene order. We initially searched for collinear groups of genes within the assembled *A. vaga* reference genome using MCScanX^22^ with E-value cut-off at ≤ 1e-05. MCScanX was run on the results of all-versus-all *blastp* search of the proteins predicted in the initial assembly. BLAST results were restricted to hits with E-value ≤ 1e-10 with the maximum number of target sequences to output per query sequence set to 5.

To focus on the genomic regions for which ploidy could be inferred with high certainty, we extracted the subset of “allelic regions” defined as collinear blocks with a high degree of collinearity (fraction of collinear genes in a block ≥ 0.7) and low synonymous divergence between haplotypes (average Ks ≤ 0.2). Coordinates of allelic regions were mapped into the coordinate system of the haploid sub-assembly, and only those regions that were fully contained within boundaries of the haploid contigs were retained. Additionally, we delineated the subset of “allelic genes” composed of collinear gene pairs embedded in allelic regions. Detailed descriptions of the procedure and of the obtained subsets are available in Supplementary Methods II.

### Mapping of Illumina reads

Trimmed Illumina 2×100 bp HiSeq reads for each sequenced individual were aligned to the initial filtered assembly and to the haploid sub-assembly with Bowtie 2 (version 2.3.2)^54^ with parameters “--no-mixed --no-discordant” and the maximum insert size of 800 base pairs. Alignments of reads to the initial assembly were used to filter out ambiguously mapped reads. Variant calling was performed from end-to-end alignments of filtered reads uniquely mapped to the haploid sub-assembly. Only properly paired reads with a high quality of mapping (MAPQ ≥ 20) were used for analyses. See Supplementary Methods III for details on filtering.

### Variant calling and filtering

Variant calls were generated using the SAMtools^55^ mpileup utility (v.1.4.1) with the parameters “-aa -u -t DP,AD,ADF,ADR” followed by the command “bcftools call” with the “-m” option. The obtained raw genotype calls were subjected to stringent filtering as follows. We excluded sites 1) within 10 bp of an indel, 2) with missing genotypes or QUAL value < 50, 3) located on contigs from the haploid sub-assembly shorter than 1,000 bp, 4) within repetitive regions, 5) with low coverage (DP<10 in any of the samples), 6) with an extremely high depth of coverage, or 7) within the windows that were outliers for SNP density. Filtering was carried out using combinations of BCFtools v.1.4.1 (https://samtools.github.io/bcftools/), VCFtools^56^ v. 0.1.15, bedtools^57^ v2.26.0, SnpSift^58^ v.4.3s utilities and awk commands. Details on filtering criteria and SNP datasets used for different analyses are provided in Supplementary Methods IV. The total numbers of sites with called genotypes in the raw and filtered SNP datasets are listed in Supplementary Table 4.

### MDS analysis

Multidimensional scaling of genome-wide identity-by-state (IBS) pairwise distances between the sequenced *A. vaga* individuals and IBS clustering was performed with PLINK^59^ v1.90b5.4. Singleton SNPs and tightly linked SNPs were excluded for this analysis. See Supplementary Methods V for details.

### Computational phasing of genotypes

Local haplotypes were assembled for each individual L1-L11 separately, using HapCUT2^16^ with the “--error_analysis_mode 1” option to compute switch error scores. For LD-related analyses, we obtained two sets of aggressively filtered phased haplotype blocks: phased dataset 1 which was used as the main dataset, and the auxiliary phased dataset 2 subjected to even more stringent filtering. From both datasets, we discarded phased blocks covered by reads supporting more than two different ‘haplotypes’ in a single individual. The logic behind this filtering step is that each individual can carry no more than two different haplotypes for a pair of SNP sites. Those pairs of sites with support for more than two ‘haplotypes’ in the aligned reads from a single individual are likely to stem from paralogous alignments and to be associated with phasing errors. Phased blocks left after this step were included in phased dataset 1. Statistics on the sizes of the resulting blocks and on the numbers of SNPs spanned by them are provided in Supplementary Tables 6 and 7. Phased dataset 2 was filtered further on the basis of presence in a block of SNPs with switch or mismatch quality values < 100. All blocks comprising more than one such SNP were discarded from phased dataset 2. Blocks harboring one such SNP were split at the corresponding site, and the chunks of the original block resulting from the split were analyzed separately.

For each individual, HapCUT2 assigns to haplotypes only those SNPs at which this individual is heterozygous. We complemented the phased haplotype blocks with the data on homozygous SNPs embedded within the phased blocks. For this purpose, we searched for homozygous SNPs flanked by heterozygous SNPs belonging to the same phased block, and assigned the homozygous SNP to both haplotypes of this block. Finally, we identified genomic segments encompassing groups of SNPs where genotypes for all individuals L4-L11, or for all individuals L1-L11, are simultaneously phased. For LD-related analyses, groups of SNPs representing different phased genomic segments were processed separately. See Supplementary Methods VI for details on filtering and processing of phased haplotype data.

### Analysis of linkage disequilibrium (LD)

To analyze patterns of LD in *A. vaga* from phased SNP data, we calculated r^2^ for each phased segment individually using VCFtools^56^ (version 0.1.15). If not stated otherwise, the reported results are based on the analysis of SNPs from the phased dataset 1. The reported results are for variants with a minor allele count of at least 4 among individuals L4-L11 or L1-L11. Results obtained with other minor allele count cut-offs were similar (data not shown). The results obtained for the more severely filtered phased dataset 2 are shown in Extended Data Fig. 2b. The rate of LD decay with physical distance was evaluated by applying LOESS regression with the smoothing parameter set to 0.4 as implemented in R (version 3.5.1). To determine the baseline r^2^ value we computed r^2^ for SNPs residing on different contigs.

See Supplementary Methods VII for details and for description of the two approaches used to infer the rate of LD decay from the unphased genotypic SNP data. Detailed explanation of the modified four-gamete test is provided in Supplementary Methods VIII.

### Signatures of recombination within individual phased genomic regions

We tested for recombination within 262 phased genomic segments of the *A. vaga* genome harboring at least 20 non-singleton SNPs simultaneously phased in all individuals L4-L11. For each such segment, we assessed whether the decay of r^2^ is significantly correlated with physical distance^17^ and performed sum of distances^18^ and pairwise homoplasy index^19^ (PHI) tests. PHI tests were performed using PhiPack and the other two types of tests using LDhat^60^. Details are given in Supplementary Methods IX.

### Construction of phylogenetic networks

Split decomposition and consensus networks were constructed and visualized with SplitsTree^21^. Description of the data used to derive consensus networks is available in Supplementary Methods X.

### Testing for Hardy-Weinberg equilibrium (HWE)

Exact tests of HWE were performed with VCFtools^56^ (version 0.1.15) on common biallelic SNPs simultaneously called in all individuals L1-L11. Tests were performed separately for the large cluster L4-L11 and for all the individuals L1-L11. To define common SNPs, we used the minor allele count cutoff of 4 in both sets of individuals. The fraction of SNPs out of HWE was computed as the fraction of common SNPs within the given set of individuals showing significant deviations from HWE (exact test P value ≤ 0.05). Details are provided in Supplementary Methods XI and Supplementary Table 8.

**Analysis of triallelic sites harboring three heterozygous genotypes and computing *H*-scores.** See Supplementary Methods XII, XIII and Supplementary Tables 9, 10 and 11.

**Estimation of the transformation rate in the *A. vaga* population.** See Supplementary Methods XIV.

**Estimation of the population-scaled recombination rate in the *A. vaga* population.** See Supplementary Methods XV.

### Identifying incongruent groupings of haplotypes and constructing phylogenetic trees

To detect incongruent groupings of haplotypes which are likely to be the result of genetic exchanges, we used the same set of 262 phased genomic segments harboring at least 20 non-singleton SNPs simultaneously phased in all individuals L4-L11 that was used to detect recombination (see section “Signatures of recombination within individual phased genomic regions”). These segments were distributed between 221 contigs belonging to the original assembly and spanned a total of 260,204 base pairs. For each such segment, we reconstructed sequences of the two haplotypes in each individual based on the corresponding sequence from the haploid sub-assembly and the set of phased non-singleton SNPs using BCFtools v.1.4.1. Only non-singleton SNPs that satisfied all filtering criteria and were simultaneously phased in L4-L11 were used to reconstruct haplotypes; the remaining sites were treated as monomorphic.

For each of the two haplotypes within each individual L4-L11, we computed the nucleotide distance (proportion of nucleotide differences) to the other haplotype within the same individual and to each haplotype in all other individuals. For each haplotype, we then identified the closest haplotypic counterpart in other individuals. To test the robustness of this matching, we compared the number of nucleotide differences between the haplotype and its closest (N1) and second closest (N2) counterpart. The haplotype was defined as having an unambiguous closest counterpart if the difference between N2 and N1 was 3 SNPs or more (N2-N1≥3).

To detect cases when the two haplotypes (H1 and H2) of a single individual clustered with haplotypes from two different individuals, we selected, for each individual, those phased segments where the closest counterpart was unambiguously identified both for H1 and H2. Among those, we searched for cases when their closest counterparts (H1′ and H2′) were found in two different individuals. Finally, we retained only reciprocal best matches. That is, we required that H1 and H2 were also identified as the closest counterparts of the haplotypes H1′ and H2′ respectively.

To exclude from consideration incongruent groupings of haplotypes that may be caused by gene conversion, we further required the distances separating pairs of identified haplotypic counterparts from different individuals (H1-H1′ and H2-H2′) to be shorter than the distances separating the two haplotypes within the same individual (for details see Supplementary Note).

In total, out of the 262 analyzed phased genomic segments, 79 exhibited incongruent groupings of the two haplotypes at least in one individual. For each such segment, we built an unrooted phylogenetic tree using maximum likelihood method in PhyML^41^ under the GTR+G model (four substitution rate categories, the gamma shape parameter estimated from the data) with 500 bootstrap trials. To check whether incongruent groupings of the two haplotypes were reliable, we parsed the resulting phylogenetic trees using ETE 3 toolkit^61^. We searched for cases when incongruent groupings of haplotypes H1-H1′ and H2-H2′ had strong bootstrap support (≥70%). In total, 56 phased segments exhibited incongruent groupings of the two haplotypes at least in one individual with strong bootstrap support.

The resulting numbers of segments with well supported incongruent groupings of haplotypes for each individual are shown in Extended Data Table 1. We also computed, for each pair of individuals, the overall number of cases where there existed a third individual harboring one haplotype clustered with a haplotype of the first individual from the pair and the other haplotype clustered with a haplotype of the second individual (Supplementary Table 12). Selected phylogenetic trees were visualized in MEGA7^62^ (root placed at the midpoint).

### Characterizing recombination in *A. vaga*

We investigated whether the probability of a recombination event is associated with the GC-content of a genomic region. As a proxy for the recombination rate of a region, we used 1) the normalized minimum number of recombination events^25^ (R_min_) inferred for individual phased segments with LDhat^60^, or 2) the fraction of SNP pairs passing the modified four-gamete test for the individual contigs from the haploid sub-assembly. To confirm that the observed negative correlation between R_min_ and GC-content is not an artifact of the larger sizes of the GC-poor segments, we performed multiple linear regression analysis with GC-content and the size of the segment as the explanatory variables and R_min_ as the response variable. To uncouple the effects of GC-content and contig size on the fractions of SNP pairs passing the modified four-gamete test for individual contigs, we compared fractions of SNP pairs passing the modified four-gamete test among SNP pairs falling within the same distance bin for contigs of different GC-content.

To see if individual recombination events tend to happen in regions with skewed GC-content, we focused on genomic intervals likely overlapping sites of true recombination events. For this purpose, for each contig from the haploid sub-assembly carrying pairs of SNPs passing the modified four-gamete test, we selected the pair of SNPs separated by the smallest distance among all recombinant SNP pairs and located further than 100 bp of each other (if there were SNP pairs satisfying these conditions). We treated intervals separating such sites as a proxy for the locations of recombination events and compared their GC-content to that of randomly sampled genomic intervals matched for number, length and contig identity.

See Supplementary Methods XVI for details and for methods used for analysis of association between recombination rate and gene features.

### Statistical analysis

Exact binomial confidence intervals demonstrated in Fig. 2f, Extended Data Fig. 4d and Extended Data Fig. 5b were computed with the function *binom.test* in R (version 3.5.1). Significance of the difference between proportions of recombinant SNP pairs for different groupings of SNP pairs meeting the conditions of the modified four-gamete test was assessed using Pearson Chi-square test with Holm correction for multiple comparisons as implemented in the *pairwise.prop.test* function in R (version 3.5.1) or two-sample Z-test for proportions (function *prop.test* in R employed without continuity correction, two-sided test).

To compare differences in distances between recombinant and non-recombinant pairs of SNPs, we employed the Mann-Whitney U (two-sided) test as implemented in the function *wilcox.test* in R (version 3.5.1).

To test for recombination in individual phased genomic segments, for each segment we computed correlation of r^2^ with distance^17^ and sum of distances between pairs of sites carrying all four haplotypes^18^ in the actual and permuted data as implemented in LDhat^60^. One-sided P values for each segment were obtained from 10,000 permutations and adjusted for multiple comparisons using the Bonferroni correction. Significance of the PHI statistic^19^ for each segment was assessed under the assumption of a normal distribution of the PHI statistic; the obtained one-sided P values were corrected by applying the Bonferroni correction.

Significance of an excess of observed over expected numbers of triallelic sites harboring all three heterozygous genotypes was assessed using one-sample Z-test for proportions (function *prop.test* in R employed without continuity correction, two-sided test).

To explore the relationship between the GC-content and recombination rate, we expressed the recombination rate as the normalized minimum number of recombination events or as the fraction of recombinant SNP pairs, calculated Pearson’s correlation coefficients (function *cor.test* in R), and performed a linear regression analysis (function *lm* in R version 3.5.1). 95% confidence bands around the regression lines in Extended Data Fig. 4a, Extended Data Fig. 5a and Extended Data Fig. 6 were obtained using the *geom_smooth* function from the package ggplot2. The average GC-content of recombinant intervals was compared to average values of GC-content in 1,000 random samples of genomic intervals matched in number, length and contig identity to the recombinant intervals; one-sided P value was obtained as the fraction of 1,000 random datasets with an average GC-content lower or equal to the average GC-content of the recombinant intervals.

## Acknowledgements

This work was partially supported by the Russian Foundation for Basic Research, research project No. 16-34-01303 mol_a to O.A.V.; MiSeq sequencing was funded by grant No. 16-14-10173 from the Russian Science Foundation to A.S.K.; Y.R.G. was supported by grant No.16-04-01579 from the Russian Foundation for Basic Research. We thank T.N. Gerasimova (Water Problems Institute of RAS) for help with species identification; Artem S. Kasianov (Skolkovo Institute of Science and Technology) for help with genome assembly and haplotype phasing; Andrei A. Minin, Arsen S. Mikaelyan and Irina Y. Bakloushinskaya at Koltzov Institute of Developmental Biology of RAS for providing access to experimental facilities, sharing reagents and experimental assistance; Sofya K. Garushyants and Elena R. Nabieva at Skolkovo Institute of Science and Technology, Dmitriy V. Vinogradov (IITP RAS) and Sergey A. Naumenko (The Hospital For Sick Children, Toronto) for comments and support.

## Author contributions

A.S.K. designed the study. A.S.K. and G.A.B. supervised and coordinated the project. Y.R.G., E.A.M. and A.S.K. performed sample collection. E.A.M. performed microscopy analysis and species identification. E.A.M. and Y.R.G. established *A. vaga* cultures. Y.R.G., S.G.O., O.A.V., I.A.Y. and E.A.M. cultivated rotifers. T.V.N., Y.R.G., O.A.V. and I.A.Y. extracted DNA. M.D.L. and A.A.P. constructed Illumina libraries and performed the sequencing. E.S.G. carried out annotation of the *A. vaga* genome. O.A.V. and G.A.B. designed the analyses. O.A.V. analyzed the data. O.A.V., G.A.B. and A.S.K. interpreted the data. A.S.K. developed equations describing how the probability of recombination depends on the transformation rate. A.O.Z. assisted in data analysis and rescued the data. I.R.A. provided advice on data analysis, edited the manuscript. G.A.B., A.S.K. and O.A.V. wrote the paper with contributions from all authors.

