## Supplementary material for "Recombination in a natural population of the bdelloid rotifer *Adineta vaga*"

**Supplementary Information for "Recombination in a natural population of the bdelloid rotifer *Adineta vaga*" by Vakhrusheva *et al.***

### Supplementary Methods

#### I. Construction of non-redundant haploid subset of the *A. vaga* genome (haploid sub-assembly)

Due to high heterozygosity, the two haplotypes of the *A. vaga* genome assemble into separate contigs at the majority of loci[1]. Still, in a substantial portion of the genome, the two haplotypes collapse into a single contig, leading to a mosaic organization of the assembly with alternating ploidy levels. This complicates application of standard variant calling and population genomics methods that presume uniform ploidy across all loci and are mainly targeted at diploid variant calls made against a haploid assembly. To overcome this difficulty, we obtained a reduced haploid representation of the *A. vaga* genome.

To retrieve a haploid subset of the assembly, we first searched for the pairs of highly similar genomic segments within the assembly likely corresponding to two haplotypes. After assigning some haplotype segments to such pairs, we retained only a single segment from a pair and discarded genomic regions without haplotypic counterparts. This procedure aims at reducing redundancy of the assembly, while simultaneously ensuring that only truly diploid loci are included into the haploid representation.

To achieve this, for each contig we identified the subset of the assembly likely containing its haplotypic counterparts. We started by carrying out all-versus-all BLAST[2] search of the filtered set of contigs within the assembly. BLAST searches were performed with *blastn* from BLAST+ 2.2.31 with the following parameters: -evalue 1e-10 -outfmt "6 qseqid sseqid pident length mismatch gapopen qstart qend sstart send evalue bitscore qlen slen" -task dc-megablast -max_hsps 1.

Next, for each contig we selected, using a custom bash script, those among the remaining contigs that had regions of high similarity to it (at least one *blastn* dc-megablast alignment with the considered contig with E-value ≤ 1e-50 and alignment percent identity ≥ 90%). We used all_bz[3] v.15 to create pairwise blastz[4] alignments between each contig and its counterparts. The initial blastz output was further processed with single_cov2 to remove secondary alignments in the contig regions where a hit to more than one other contig region was found. This resulted, for each contig, in a set of pairwise alignments such that each position of the contig was aligned to no more than one position of the matched contigs. Thus, the resulting set of alignments for a contig could be viewed as the set of best BLAST hits between it and the rest of the assembly.

To extract only those pairs of genomic segments that are reciprocal best matches, we devised a procedure analogous to identifying reciprocal best BLAST hits among genes. For each alignment between contig A and contig B belonging to the set of best matching alignments of contig A (let us denote it by ‘forward alignment’), we determine if there is a corresponding alignment between contig B and contig A among the set of best matching alignments of contig B (‘reverse alignment’). We retain the pair of the aligned segments if the coordinates of the forward and reverse alignments are identical or if their shared span covers ≥ 80% of the longer alignment. In the latter case, the boundaries of the reciprocally best matching segments were defined as the overlap between the forward and reverse alignments.

The resulting set of reciprocally best matching genomic segments is likely to represent pairs of haplotypes. To obtain a non-redundant haploid subset of the assembly, from each such pair of best matching segments, we selected the one from the longer contig, and discarded the remaining one. To increase continuity of the haploid sub-assembly, if a pair of non-redundant haploid segments was located on the same contig and separated by ≤ 200 base pairs, we took a union of such segments. The statistics for the resulting haploid sub-assembly are shown in Supplementary Table 2. The above-described steps were carried out with a series of custom Perl scripts.

The final haploid sub-assembly spanned 76,679,421 base pairs (Supplementary Table 2), suggesting that at least ~80% of the initial assembly is represented by two haplotypes.

### II. Identification of allelic regions and allelic genes

To test the robustness of our results against erroneous identification of haplotype pairs, we separately analyzed the fraction of the haploid sub-assembly covered by long blocks of genes that are collinear between the two haplotypes in the diploid genome.

To obtain a subset of the *A. vaga* genome with high-confidence ploidy, we identified genomic regions that could be assigned into pairs of highly similar segments with conserved gene order. We initially searched for collinear groups of genes within the assembled *A. vaga* reference genome. For this, we first ran an all-versus-all *blastp* search of the proteins predicted in the *A. vaga* genome (carried out with BLAST+ 2.2.31). BLAST results were restricted to hits with E-value ≤ 1e-10 with the maximum number of target sequences to output per query sequence set to 5, and self-to-self hits were discarded. Next, to identify collinear groups of genes, we ran MCScanX[5] on the output of *blastp* with an E-value cut-off of ≤ 1e-05. This resulted in a total of 1,770 detected syntenic blocks.

Next, for each syntenic block, we calculated the fraction of collinear genes and the average value of Ks across all pairs of collinear genes using a custom Perl script. Numbers of synonymous substitutions per site (Ks) for each matched pair of collinear genes within the block were computed using the script add_ka_and_ks_to_collinearity.pl distributed as a part of the MCScanX package. The fraction of collinear genes for a block was computed as the number of collinear gene pairs divided by the maximum number of genes between two genomic regions forming a collinear block.

Presence of gene duplications might inflate the total number of genes covered by a collinear block and cause a downward bias in the estimated fractions of collinear genes. To account for this, we subtracted the total number of tandem duplications identified by MCScanX within each genomic region from the total number of genes covered by the considered region.

Distribution of average Ks values per collinear block versus fractions of collinear genes revealed two clearly distinguishable groups of blocks (Fig. 1b). The group with high fractions of collinear genes and low average values of Ks (Fig. 1b, blue dots) is likely to correspond to pairs of ancestral haplotypes, while the other group exhibiting lower extent of collinearity and higher synonymous divergence (Fig. 1b, red dots) most probably stems from ancient whole-genome duplication. Such genome organization confirms the same pattern that has already been reported for the first published genome of *A. vaga*[1].

To focus on the genomic regions for which the ploidy could be inferred with high certainty, we extracted a subset of the collinear blocks with a high degree of collinearity and low synonymous divergence (hereafter referred to as "allelic regions"). These are the regions that are most likely to be represented in the assembly by two haplotypes. We delineate the allelic regions as a subset of genomic segments possesing a within-genome counterpart with a high fraction of collinear genes (fraction of collinear genes in a block $\geq 0.7$) and low average values of Ks (Ks $\leq 0.2$) (Fig. 1b). A total of 1,387 collinear blocks satisfied these criteria, with 1,354 blocks remaining after removal of the conflicting synteny segments encompassing overlapping genomic regions. In addition to the subset of allelic regions, we specify the subset of the genes embedded in allelic regions (hereafter referred to as "allelic genes"). The initial subset of allelic genes was composed of 12,489 collinear gene pairs residing within the allelic regions and filtered for the individual values of Ks (Ks $\leq 0.2$).

To delineate haploid equivalent of the allelic regions, for each of the 1,354 pairs of collinear allelic segments, we left only a single segment, retaining the one from a longer contig in a pair and discarding its counterpart. This non-redundant subset of unique non-overlapping allelic regions spanned 34,691,452 base pairs.

Having thus obtained a non-redundant haploid representation of regions with high-confidence ploidy, we mapped it into the coordinate system of the original haploid segments identified in the previous step (see section «Construction of non-redundant haploid subset of the *A. vaga* genome»). We retained only those allelic segments that were fully contained within boundaries of the original haploid segments; those with partial overlaps were discarded. A total of 833 allelic segments spanning 19,300,566 base pairs and encompassing 7,245 allelic genes remained after this step. We used these final subregions of the haploid segments throughout the paper as portions of the genome with high-confidence ploidy. The designations "allelic regions" and "allelic genes" in the main text of the paper refer to these final sets of 833 regions and 7,245 genes respectively.

The procedures described above were carried out with custom Perl scripts.

### III. Mapping of Illumina reads

We aligned Illumina 2×100 bp HiSeq reads generated for each sequenced individual to:

1) Initial filtered diploid contigs.

2) Non-redundant haploid segments (see section «Construction of non-redundant haploid subset of the *A. vaga* genome»).

Alignments of reads to the diploid contigs were used to filter out ambiguously mapping reads, prior to performing the alignment of reads to the haploid sub-assembly. Actual identification of variable sites was performed on read alignments to the haploid sub-assembly, which allowed us to assume diploid samples during variant calling.

We mapped reads with Bowtie 2 (version 2.3.2)[6]. The choice of this aligner was motivated by its ability to find global end-to-end alignments of reads to the reference genome. This is advantageous compared to local read alignment when dealing with the genome rich in repetitive sequences as is the case with genomes of bdelloid rotifers bearing remnants of a whole-genome duplication[1].

First, we mapped trimmed reads from each individual to the original filtered contigs using Bowtie 2 with the parameters "--no-mixed --no-discordant" setting the maximum insert size to 800 base pairs and allowing to report up to 5 distinct alignments. The overall alignment rates for different individuals ranged from 74.29% to 93.43%.

Assuming that most genomic loci in the assembly are represented by two haplotypes, we would expect no more than two 'true' alignments per a pair of reads. Those reads mapping to more than two genomic locations are likely to produce spurious alignments to paralogous regions.

To avoid using erroneous alignments for variant identification, which could result in false positive variant calls, we removed reads mapping to more than two loci in the diploid genome prior to aligning the reads to the haploid sub-assembly. For this purpose, for each pair of reads, we tabulated the number of properly paired alignments in the diploid genome, leaving only reads that were mapped in a proper pair to one or two locations. This filtering step was accomplished using seqtk v.1.2-r94 (https://github.com/lh3/seqtk) and a combination of custom scripts written in Perl and bash.

For each sequenced individual, subsets of reads that passed this filtering step were in turn remapped to the non-redundant haploid segments. Alignment of the filtered subsets of reads against the haploid sub-assembly was performed with Bowtie 2 with the parameters "--no-mixed --no-discordant" specifying the maximum insert size of 800 base pairs with only a single best alignment of the pair of reads reported.

To further reduce the number of erroneous mappings, we parsed SAM files with the reads mapped against the haploid sub-assembly using a custom Perl script and removed alignments of reads for which more than one valid alignment in the haploid genome was found (those with XS tag set). We filtered out alignments of both paired-end reads, irrespective of whether a secondary alignment was found for a single read or for both reads forming the pair.

We used SAMtools[7] v.1.4.1 (http://samtools.sourceforge.net/) to convert the filtered SAM files to sorted BAM files and perform additional filtering on the mapping quality (MAPQ) retaining only those reads that have MAPQ $\geq 20$. The resulting BAM files with paired-end alignments left after the above-described filtering steps were used for variant calling.

### IV. Variant calling and filtering

Throughout the analyses, we use two genotypic data sets. SNP dataset I includes SNP calls for variable sites segregating among the individuals L1-L11. SNP dataset II comprises calls for both variable and invariant sites.

A stringently filtered subset of the SNP dataset I was used for local haplotype reconstruction via read-based phasing. We devised SNP filtering approach for this dataset in such a way as to maximally reduce the percentage of false positive variant calls which can potentially lead to phasing errors. For this, prior to performing read-based phasing, we removed from the SNP dataset I all sites with more than two nucleotides present in the aligned reads, even if some of the nucleotides did not occur in any of the genotypes. For these purposes, all nucleotides present in the aligned reads were treated as putative alleles in the process of variant calling, regardless of whether they were supported by any of the called genotypes.

Conversely, the SNP dataset II was primarily intended for a survey of triallelic SNPs and computing the pairwise genotypic distances. Treating all nucleotides at a particular site present in reads but not appearing in the resulting genotypes as alleles might lead to erroneous classification of sites with respect to the number of segregating alleles. Therefore, for the purposes of building the SNP dataset II, to minimize the number of sites misidentified as being triallelic due to sequencing errors, only those nucleotides at a given site supported by at least one of the genotypes were regarded as alleles.

Single-nucleotide variants were called from read alignments to the haploid sub-assembly. As a result, we were calling diploid variants, because homologous sites of both haplotypes were aligned to the same site of the sub-assembly. Genotype calls in both datasets were generated using the SAMtools[7] mpileup utility (v.1.4.1) with the parameters "-aa -u -t DP,AD,ADF,ADR" followed by the command "bcftools call" with the "-m" option. To identify all alleles present in the reads, including those potentially absent from called genotypes, and to skip invariant sites, "bcftools call" was run with the additional parameters "-A" and "-v". These additional parameters were employed to produce genotype calls included in the SNP dataset I.

Next, we performed stringent filtering of the obtained raw genotype calls. We successively applied a series of filters, removing sites falling into one or more of the following categories from the datasets:

1) Sites residing within 10 bp of an indel.
2) Sites with missing genotypes or QUAL value < 50.
3) Sites located on haploid segments shorter than 1,000 bp.
4) Sites residing in repetitive regions.*
5) Sites with low coverage (DP<10 in any of the samples).
6) Sites with extremely high depth of coverage.**
7) Sites residing within the windows outliers for SNP density.***

*Annotation of repetitive regions in the haploid sub-assembly of the *A. vaga* genome was carried out with RepeatMasker (version open-4.0.7) (http://www.repeatmasker.org/).

**For SNP dataset I, which includes only variable sites, we removed from further consideration sites identified as being outliers with respect to the mean coverage across 11 individuals, the total coverage summed across 11 individuals or individual coverage values as determined for each sample separately. Identification of outliers was performed using the interquartile range method in R (version 3.3.2). For SNP dataset II, we discarded all sites with depth of coverage DP > 300 in any of the individuals.

***Genomic regions with unusually high densities of variable positions are likely to stem from reads mapping to paralogous regions. To avoid using false positive variant calls resulting from misalignment of such reads, we searched for outlier regions with respect to SNP density and discarded variants falling within such regions. For this purpose, we conducted a sliding window analysis (using a window length of 1,000 bp and a step size of 500 bp) of the *A. vaga* haploid sub-assembly, computing fractions of segregating sites based on the SNP dataset I in each window. Detection of outliers was performed using the interquartile range method in R (version 3.3.2).

Filtering was carried out using combinations of BCFtools v.1.4.1 (https://samtools.github.io/bcftools/), VCFtools[8] v. 0.1.15, bedtools[9] v2.26.0, SnpSift[10] v.4.3s utilities and awk commands.

The total numbers of sites with called genotypes in the raw SNP datasets and the numbers of sites remaining after successive application of various filters are listed in Supplementary Table 4. The final subsets of the SNP dataset I and II that passed all the filters are further referred to as the stringent datasets I and II respectively.

### V. MDS analysis and identification of the population outliers

We performed multidimensional scaling (MDS) analysis of genome-wide identity-by-state (IBS) pairwise distances between the sequenced *A. vaga* individuals with PLINK[11] v1.90b5.4.

Variant calls from the stringent SNP dataset I were additionally filtered prior to processing with PLINK. A list of variants was thinned in such a way that the resulting dataset did not contain any variants within 10 bp of one another. We also excluded from consideration singleton variants and tightly linked variants (defined as pairs of variants residing within 1,000 bp of each other with $r^2$ values greater than 0.3). The resulting subset of SNPs was retained for the MDS analysis.

Visual inspection of MDS plots revealed partitioning of the sequenced individuals into two groups, with one group formed by individuals L1-L3 and another one by individuals L4-L11 (Fig. 1c). To detect potential population outliers, for each individual we determined five closest neighbors in terms of identity-by-state distances. However, none of the between-individual comparisons showed significantly negative Z-scores (below -3), revealing no outliers that could be identified in this way.

Although no outliers were detected using the Z-score method, IBS clustering analysis provided statistical evidence for the presence of at least two genetically distinct groups among the sequenced individuals. IBS clustering analysis was carried out with varying thresholds on P values used to discriminate clusters. While the number of detected clusters varied from 2 for P value set to 1e-100 to 5 for P value set to 1e-3, three individuals (L1, L2 and L3) including the reference one (L1) were always ascribed to a separate cluster, hereafter referred to as the small cluster. These three individuals were also found to be outliers in terms of fractions of identity-by-decent (IBD) SNPs (according to interquartile range method of outlier detection) with respect to the rest of the sample, thus providing statistical justification for separate analysis of individuals L4-L11 (hereafter referred to as the large cluster).

The average pairwise genotypic distance was 1.56% for the individuals belonging to different clusters, 0.85% for the 3 individuals belonging to the small cluster, and 0.67% for the 8 individuals belonging to the large cluster (Supplementary Table 5). Out of the 11 individuals used in the study, nine (L1-L4 and L6-L10) were sampled from the Moscow region and two, L5 and L11, sampled from the Kostroma region, 550 km to the NE. Despite this distance between the two sampling locations,

L5 and L11 clearly belong to the large cluster, together with individuals L4 and L6-L10.

To minimize the potential effect of population structure, we focused the subsequent analyses on the large cluster. If not indicated otherwise, the reported results are based on the analysis of this cluster. Nonetheless, the main findings of the manuscript were recapitulated on the complete set of the sequenced individuals (L1-L11) (see below; Extended Data Fig. 2c; Extended Data Fig. 3b, c; Supplementary Table 8; Supplementary Table 9), confirming the robustness of our conclusions to the effects of population stratification.

### VI. Computational phasing of genotypes

We performed computational phasing of genotypes using biallelic SNPs from the stringent SNP dataset I (n = 1,774,991) and the strictly filtered alignments of reads to the haploid sub-assembly (see sections «Mapping of Illumina reads» and «Variant calling and filtering»). To mitigate the impact of sequencing errors on haplotype reconstruction, we applied further stringent criteria for inclusion of SNPs in the dataset subjected to phasing, discarding all sites with more than two different nucleotides present in the aligned reads across the individuals L1-L11. Local haplotypes were assembled for each sample L1-L11 individually, using HapCUT2[12] with the "--error_analysis_mode 1" option to compute switch error scores.

Phased haplotype blocks were aggressively filtered before being used for subsequent analyses. The logic behind the main filtering step is that each individual can carry no more than two different haplotypes for a pair of SNP sites. Those pairs of sites with support for more than two 'haplotypes' in the aligned reads from a single individual are likely to stem from paralogous alignments and to be associated with phasing errors. We discarded phased blocks encompassing such sites prior to the analysis, as their presence might create artifactual evidence for LD decay.

For this purpose, we parsed fragment matrix files generated by HapCUT2 for each individual, L1-L11, and extracted information on the haplotypes supported by reads for each pair of the SNPs phased in a given individual. We designated pairs of SNPs represented by more than two 'haplotypes' in the aligned reads from a single sample as conflicting. Most such cases originated from sequencing errors with the third (least frequent) 'haplotype' supported only by a single read. Thus, we narrowed the list of the conflicting SNP pairs down to those present in reads as three distinct haplotypes each supported at least by two reads. We also regarded as 'conflicting' all SNP pairs represented by four haplotypes in a single individual irrespective of the number of reads supporting different 'haplotype' variants. Having obtained lists of 'conflicting' SNP pairs for each individual, we removed phased blocks encompassing such SNPs from further consideration.

Phased blocks remaining after applying this filter were used as the main phased dataset (further referred to as 'phased dataset 1'). Statistics on the lengths of phased blocks included in the phased dataset 1 for different individuals and on the numbers of variants spanned by such blocks are provided in Supplementary Table 6 and Supplementary Table 7.

To see whether the patterns in LD decay depend on the stringency of filtering criteria and to show that it cannot be explained solely by phasing errors, we also obtained a more strictly filtered phased dataset ('phased dataset 2'). For this purpose, in addition to removing phased blocks with 'conflicting' pairs of SNPs, we subjected sets of locally phased haplotypes to further filtering. We used phred-scaled probabilities of switch errors and mismatches generated by HapCUT2. For each phased block left after removal of blocks with ‘conflicting’ pairs of SNPs, we considered SNPs with values of switch or mismatch quality < 100 as problematic. All blocks comprising more than one problematic SNP were completely discarded from the dataset. Blocks with a single problematic SNP were split at the corresponding site, and the chunks of the original block resulting from the split were analyzed separately. Detection of ‘conflicting’ SNP pairs and subsequent filtering of HapCUT2 output files was conducted with custom Perl scripts. For both filtered datasets, the resulting files with the phased blocks in the HapCUT2 format were converted to VCF format using the utility HapCutToVcf from fgbio (http://fulcrumgenomics.github.io/fgbio/).

For each individual, HapCUT2 assigns to haplotypes only those SNPs at which that individual is heterozygous; consequently, all homozygous sites are omitted from the output. However, sites that are in a homozygous state in some individuals may occur in a heterozygous state in other individuals. Therefore, data on the homozygous sites are essential when exploring haplotypic data across several individuals simultaneously. To complement the phased haplotype blocks with the data on SNPs at which a given individual is homozygous, we searched for cases where a homozygous site is embedded within a phased block. For this, for each homozygous site, we identified closest flanking SNPs at which the individual in question is heterozygous. We regarded a homozygous SNP as embedded in a phased block, if both its left and right closest heterozygous SNPs were phased and belonged to the same phased block. In this case, we assigned the homozygous SNP to the block encompassing its heterozygous neighbors, assuming that both haplotypes carry the same variant.

After adding ‘phasing’ information for the homozygous variants, we further processed the VCF files and identified haplotype blocks nested within other blocks, removing such cases from the analysis. The above-described processing procedures applied to the phased VCF files were carried out with a series of custom Perl scripts, if not stated otherwise.

Next, we identified genomic segments encompassing groups of variable sites where genotypes for all the individuals L4-L11, or for all the individuals L1-L11 are simultaneously phased. In the text of the paper, we refer to such genomic segments harboring at least two sites simultaneously phased in L4-L11 or in L1-L11 as phased genomic segments. For each such phased genomic segment, we extracted the corresponding portion of the VCF file into a separate VCF file with the aid of awk commands. We also obtained subsets of the variants belonging to individual phased genomic segments applying different thresholds on a minor allele count among individuals L4-L11 or L1-L11 using BCFtools v.1.4.1 (https://samtools.github.io/bcftools/).

For the purposes of calculating $r^2$ values and other LD-related analyses, groups of variants representing different phased genomic segments were processed separately.

### VII. Analysis of linkage disequilibrium (LD)

We calculated $r^2$ for each phased segment individually from phased SNP data using VCFtools[8] (version 0.1.15). If not stated otherwise, the reported results are based on the analysis of the SNPs from the phased dataset 1. The reported results are for variants with a minor allele count of at least 4 among individuals L4-L11 or L1-L11. Results obtained with other minor allele count cut-offs were similar (data not shown). The results obtained for the more severely filtered phased dataset 2 were also qualitatively similar (Extended Data Fig. 2b). We also recapitulated the main findings on the subset of those SNPs from the phased dataset 1 that are covered by long allelic regions (Extended Data Fig. 2a).

To determine the baseline $r^2$ value, we computed $r^2$ for sites residing on different contigs in the original assembly. If the total number of site pairs from different contigs in the dataset exceeded 10,000,000, we thinned the dataset by randomly drawing 10,000,000 pairs of sites. In this case, the displayed distributions of inter-contig $r^2$ values and the corresponding mean and median $r^2$ values are based on the thinned datasets.

The rate of LD decay with physical distance expressed in base pairs was evaluated with LOESS regression with the smoothing parameter set to 0.4 as implemented in R (version 3.5.1). We also estimated the rate of short-range decay of $r^2$ with physical distance by applying a nonlinear regression model assuming recombination-drift equilibrium (see section «Estimation of the population-scaled recombination rate»).

To make sure that the observed LD decay was not an artifact of phasing, we assessed LD decay directly from the unphased genotypic SNP data by using two approaches. The first approach is based on inferring haplotypes on the basis of variable homozygous sites. The rationale behind is as follows. For each individual, it is possible to determine haplotypes for sites at which this individual is homozygous, as phase of homozygous SNPs on the same contig is already "known". We make use of this by comparing haplotypes of variable sites at which each individual is nonetheless homozygous. For this purpose, we selected biallelic sites variable among individuals L4-L11 such that each individual is homozygous at each site (n = 18,995). We further filtered out sites with the least frequent genotype private to a single individual, leaving only those sites where each of two homozygous genotypes (0/0 and 1/1) was present at least in two individuals (n = 3,410). This requirement automatically filters out variants with minor allele count (MAC) below 4. To retain only truly homozygous genotypes, we excluded all sites at which more than one nucleotide occurred in reads in any single individual. We also required genotypes to be simultaneously supported by forward and reverse reads in all individuals. These filters resulted in the final set of 2,573 variable sites. We converted GT field of such sites in the VCF file to the format of a phased genotype (0/0 -> 0|0; 1/1 -> 1|1) and used the resulting VCF file containing 2,573 sites to compute $r^2$ values with VCFtools (Fig. 2b).

The second approach to inferring the rate of LD decay from the unphased genotypic SNP data relies on calculation of squared correlation coefficients between genotypes. Squared correlation coefficients were computed for comparisons of 10,000 randomly drawn biallelic sites versus the rest of the segregating biallelic sites using VCFtools. SNP pairs were binned according to the distance separating the pair at resolution of 10 base pairs and average squared correlation coefficient between genotypes was determined for each bin. Confidence intervals for average genotypic correlation coefficient at different distances were derived from 1,000 bootstrap samples. Only bins with SNPs at a distance of ≤ 200 base pairs are shown in Fig. 2c, as there was almost no further decrease in the average squared correlation coefficient between genotypes after this distance. Fig. 2c shows the plot for whole-genome variants; the variants belonging to the genomic segments with high-confidence ploidy recapitulated the same pattern of LD decline (not shown).

### VIII. Distinguishing signatures of recombination and of gene conversion

To disentangle signatures of recombination from those of gene conversion, we devised a modified implementation of the Hudson's four-gamete test[13]. In the original Hudson's four-gamete test, the presence of all four possible haplotypes for a pair of biallelic polymorphic loci within a population is interpreted as evidence for recombination, because recurrent mutations are unlikely. However, a mutation followed by gene conversion within a single individual would suffice to explain the presence of all four haplotypes without assuming genetic exchanges between individuals (Fig. 2d). Nevertheless, allelic gene conversion can only produce a homozygous genotype from the heterozygous one, but not *vice versa*. Therefore, it cannot produce a pair of individuals, each heterozygous at two loci, carrying all four haplotypes (Fig. 2e); while such a pair can obviously arise through recombination involving genetic exchanges. We use this feature of gene conversion to distinguish it from genetic exchanges.

For this purpose, for each pair of the sequenced *A. vaga* individuals, we consider only those pairs of sites at which both individuals are simultaneously heterozygous. Next, among all such pairs of heterozygous sites for a given pair of individuals, we look for those that are represented by all four possible haplotypes in these two individuals. In the absence of recurrent mutations, presence of such pairs of SNPs is indicative of recombination. We refer to such pairs of sites in the text of the paper as to 'recombinant' pairs of sites, or pairs of sites passing the modified four-gamete test.

To obtain a statistic that could be applied to all individuals simultaneously, we compute the fraction of SNP pairs passing the modified four-gamete test among all SNP pairs that are simultaneously heterozygous in at least one pair of the considered individuals.

To see if the fraction of recombinant SNP pairs increases with the distance, heterozygous SNP pairs meeting the requirements of the modified four-gamete test were subdivided into 4 distance bins with approximately equal numbers of cases using the *cut_number* function from the package ggplot2 in R version 3.5.1. Proportions of recombinant SNP pairs were calculated for each bin, and significance of the difference between proportions for all pairs of bins was assessed using Pearson Chi-square test with Holm correction for multiple comparisons as implemented in the *pairwise.prop.test* function in R (Fig. 2f). To complement this analysis, we compared distributions of distances between recombinant and non-recombinant pairs of SNPs meeting the conditions of the modified four-gamete test showing that recombinant pairs of SNPs tend to reside farther apart from each other than non-recombinant ones (Fig. 2g; Mann-Whitney U test, $P < 2.2e-16$).

The observed increase in the fraction of recombinant SNP pairs with increasing physical distance (Fig. 2f, g) is equivalent to LD decay that could not be ascribed solely to action of gene conversion. The results reported in the paper are for pairwise comparisons among individuals L4-L11 with the cut-off threshold for a minimal allele count of 4. The results obtained using the complete set of individuals, L1-L11, or different cut-offs on a minimal allele count showed the same general trend (data not shown).

### IX. Signatures of recombination within individual phased genomic regions

We explored signatures of recombination within the individual phased genomic regions applying two permutation tests to the segments of the *A. vaga* genome harboring at least 20 non-singleton SNPs simultaneously phased in all the L4-L11 individuals. A total of 262 segments that satisfy these conditions were distributed between 221 contigs belonging to the original assembly.

First, for each such segment we assessed whether the decay of $r^2$ is significantly correlated with physical distance[14]. Then, we performed the sum of distances test[15], assessing whether the sum of distances between segregating sites harboring all four possible haplotypes is significantly larger than that expected by chance based on the value of the statistic in the permuted data. Both tests were carried out using LDhat[16], and a one-sided P value for each considered segment was obtained from 10,000 permutations.

We also tested for recombination applying pairwise homoplasy index[17] (PHI) test (as implemented in the PhiPack) to the same set of 262 segments. The window size for computing PHI statistic was set to 100 nucleotides, and significance was assessed under the assumption of normal distribution of the PHI statistic. Split decomposition networks of the selected genomic segments for which results of PHI-test remained significant after applying the Bonferroni correction were built and visualized with SplitsTree[18]. Out of the 262 segments, 230 demonstrated significant negative correlation of $r^2$ with the physical distance at the 0.05 significance level (with 133 remaining significant after correcting for multiple testing).

The sum of the distances and PHI-tests directly assessing whether genealogical incongruence between polymorphic sites increases with physical distance also provided evidence for recombination. 207 and 235 segments out of 262 showed evidence for recombination at the 0.05 significance level according to the sum of the distances and the PHI-test, respectively, with the results for 95 and 167 segments remaining significant after the Bonferroni correction. Contradictory groupings of different individuals produced by different sets of variable sites are visualized through split decomposition networks constructed for the individual phased genomic segments (Fig. 3d).

### X. Reconstruction of consensus networks

We reconstructed consensus networks based on the collections of neighbor-joining trees built for individual regions of the *A. vaga* genome with high-confidence ploidy. Neighbor-joining trees and consensus networks were built using the program SplitsTree4[18]. Networks include splits that appear in at least 10% of the original trees.

Consensus networks were obtained for two different types of genomic loci:

a) allelic regions (regions of the haploid sub-assembly covered by long collinear blocks of highly similar genes, see section «Identification of allelic regions and allelic genes»)

b) allelic genes (genes with an allelic counterpart residing within the allelic regions)

First, we separately identified subsets of such loci carrying considerable numbers of variable sites (as determined based on the stringent SNP dataset I) within the large cluster (L4-L11) and among all individuals (L1-L11). In both cases, we used 100 as a cutoff for the minimal number of biallelic SNPs (from the stringent SNP dataset I) harbored by an allelic region, which resulted in 796 and 832 allelic regions for individuals L4-L11 and L1-L11, respectively.

For the allelic genes, we used a cutoff value of 50 for the minimal number of biallelic sites (from the stringent SNP dataset I) variable among individuals L4-L11 and a cutoff value of 100 for individuals L1-L11 and identified, respectively, 680 and 634 allelic genes meeting these requirements.

For each of the four resulting sets of the genomic segments we generated a collection of neighbor-joining trees, which were used for subsequent construction of consensus networks. Neighbor-joining trees for the individual genomic segments were built using matrices of pairwise distances between the considered individuals as input.

Distance matrices used for tree construction were computed from the unphased genotype data. The distance between a pair of individuals at a given genomic locus with $n$ polymorphic sites was calculated as a sum of genotypic distances for each polymorphic site belonging to the locus normalized by $2n$. The pairwise genotypic distance for a given polymorphic site is defined as the difference between the harbored numbers of non-reference variants. For example, the distance between the genotype 0/1 and the genotype 0/0 is 1, and the distance between genotypes 0/0 and 1/1 is 2.

### XI. Testing for Hardy-Weinberg equilibrium

Exact tests of HWE were performed with VCFtools[8] (version 0.1.15) on common biallelic SNPs (belonging to the stringent SNP dataset II) simultaneously called in all individuals L1-L11. Tests were performed separately for the large cluster L4-L11 and for all individuals L1-L11. To define common SNPs we used a minor allele count cutoff of 4 in both sets of individuals. The fraction of SNPs out of HWE was computed as the fraction of common SNPs within the given set of individuals showing significant deviations from HWE (exact test P value $\leq 0.05$).

Observed-to-expected ratios for the numbers of homozygous and heterozygous genotypes were computed for the sites with unambiguously identified major and minor alleles. To distinguish between major and minor alleles, we required the major allele count to be at least 10 among the individuals L4-L11 or 12 among the individuals L1-L11. We computed observed-to-expected ratios for the numbers of genotypes homozygous for the major allele as well as for the numbers of heterozygous genotypes. The values expected under HWE were obtained with VCFtools. Fractional expected genotype counts not ending in 0.5 were rounded to the nearest integer number.

We performed this analysis for whole-genome variants as well as for subsets of the variants residing within the regions of the *A. vaga* genome with high-confidence ploidy (allelic regions and allelic genes). The overall numbers of analyzed SNPs, fractions of SNPs out of HWE and statistics on the observed-to-expected ratios for the numbers of homozygous and heterozygous genotypes for each dataset are provided in Supplementary Table 8.

### XII. Identification of sites harboring three heterozygous genotypes

To estimate the observed to expected ratio of the numbers of triallelic sites carrying all three heterozygous genotypes, we used only those sites of the *A. vaga* genome (belonging to the stringent SNP dataset II) that were simultaneously called in all the individuals (L1-L11) and applied additional strict filters on the SNP quality. Prior to the analysis, we excluded all sites for which there were more than two nucleotides simultaneously present in the aligned reads in any individual genome. We subdivided the resulting set of sites according to the number of alleles they carried within the large cluster (L4-L11) or among all individuals (L1-L11).

The probability of a mutation recurrently affecting the same site could be estimated from the fraction of triallelic sites among all sites with two or three alleles. Hence, we calculated the fraction of triallelic sites (P3) among all sites represented by two or three alleles. This fraction could be viewed as an estimate of a probability of a mutation recurrently affecting the same site in the history of the sample of genotypes. Therefore, the expected number of triallelic sites simultaneously harboring all three possible heterozygous genotypes due to recurrent mutations could be estimated as N3*P3, where N3 is the observed number of the triallelic sites. The significance of the enrichment in the number of sites with all three heterozygous genotypes was assessed with one-sample Z-test for proportions (function *prop.test* in R employed without continuity correction).

Among high-quality 1,136,041 sites variable among the individuals L4-L11, 9,738 sites (0.0086) were found to be triallelic, thus we would expect 83.5 (9,738*0.0086) triallelic sites to carry all the three possible heterozygous combinations of alleles due to recurrent mutations. However, the observed number of such sites is 1,839 (0.189 among all triallelic sites). To explain this observation under the hypothesis of obligate asexuality, one would have to allow the rate of recurrent mutations to be ~22 times higher than it apparently is ($P < 2.2e-16$, one-sample Z-test for proportions).

Besides, recurrent mutations affecting triallelic sites should give rise to the similar numbers of tetraallelic sites and sites carrying all three heterozygous genotypes. Therefore, the observed number of tetraallelic sites could be used to obtain an independent estimate of the expected number of sites represented by three heterozygous genotypes due to recurrent mutations.

Although on average a recurrent mutation at a triallelic site is only twice more likely to produce a site with three heterozygotes than a tetraallelic site, only 1 high-quality tetraallelic site was identified among the whole-genome calls for L4-L11 individuals (Supplementary Table 9). The minimal observed ratio of sites carrying three heterozygotes to tetraallelic sites among the analyzed datasets was 511 (whole-genome calls, individuals L1-L11; Supplementary Table 9), strongly arguing against recurrent mutations as the main source of sites harboring three heterozygous genotypes. Importantly, if only the regions of the genome with high-confidence ploidy (allelic regions or allelic genes) were considered, even a greater enrichment with triallelic sites carrying three heterozygous genotypes was observed (Supplementary Table 9), making erroneous read mappings an unlikely explanation for the phenomenon.

Intriguingly, the observed to expected ratios for the numbers of triallelic sites represented by three heterozygotes were higher when the individuals L4-L11 forming the large cluster were analyzed separately than when all the individuals L1-L11 were analyzed together (Supplementary Table 9). The observed to expected ratios ranged from 12.6 to 13.3 for the individuals L1-L11 and from 22.0 to 28.1 for the individuals L4-L11 (Supplementary Table 9). More frequent genetic exchanges between genetically more similar individuals could explain this difference.

### XIII. Analysis of sites harboring three heterozygous genotypes and computing *H*-scores

It is conceivable that sites with all three possible heterozygous genotypes could simply be a result of cross-sample contamination. However, if it were the case, we would expect those samples that originated from contamination to harbor the majority of rare heterozygous genotypes. To confirm that the sites carrying all three possible heterozygous genotypes are not likely to be due to cross-sample contamination, we separately considered those sites carrying all three heterozygotes among the individuals L4-L11 that harbor only one private heterozygous genotype (n = 607). That is, we retained a site for the analysis if the least frequent of the three heterozygous genotypes was present in a single individual with the next frequent genotype present at least in two individuals.

Such private heterozygous genotypes possessed by a single individual are most likely to stem from contamination. Moreover, should contamination be the case, we would expect to see a skewed distribution of per individual numbers of such private heterozygous sites with the samples resulting from contamination carrying disproportionally more private heterozygotes.

Following this logic, we analyzed how the 607 private heterozygous genotypes are distributed among different individuals. For this purpose, for each individual, we tabulated the total number of sites with the least frequent heterozygous genotype private to this individual.

Contrary to what would be expected under contamination, we observed that the resulting numbers of private heterozygous sites were very similar across different individuals (average number of private heterozygous sites per individual was 75.875, with the minimal and maximal values of 64 and 88 sites respectively; Supplementary Table 10). Thus, the distribution of unique heterozygous genotypes among the sequenced individuals argues against contamination being the source of sites harboring all three heterozygotes.

Using the same line of reasoning, one can evaluate, for each pair of individuals, how likely it is to observe a site harboring all three heterozygotes as a result of hypothetical contamination or exchange between these two individuals. For this purpose, we computed pairwise scores (hereinafter referred to as scores ***H***) reflecting the relative probability for a given pair of individuals to have contributed to the pool of sites carrying all three heterozygous genotypes.

Let us consider a single triallelic site represented by three heterozygous genotypes A1/A2, A1/A3 and A2/A3 among the individuals L4-L11, where A1, A2 and A3 denote the three segregating nucleotides. Here we assume that A2/A3 is the only private heterozygote among the three, with the other two harbored by at least two individuals each. If the heterozygote A2/A3 resulted from a hypothetical contamination involving a pair of the sequenced individuals, then there is a limited set of the genotypes that could have given rise to this combination of alleles. Namely, the A2 allele could have been contributed by the individuals carrying genotypes A1/A2 or A2/A2, while the A3 allele could have come from the genotypes A1/A3 or A3/A3.

Therefore, the probability for a given individual K to have contributed allele A2 to the private heterozygous genotype A2/A3 could be computed as:

$$\boldsymbol{P2(K)} = \frac{\boldsymbol{W2_K}}{\sum_{I=1}^{N-1} \boldsymbol{W2_I}}$$

where

$$\boldsymbol{W2_X} = \begin{cases} \mathbf{2}, \ \textit{if the individual X is homozygous for the allele A}2 \\ \mathbf{1}, \ \textit{if the individual X is heterozygous for the allele A}2 \\ \mathbf{0}, \ \textit{otherwise} \end{cases}$$

and summation $\sum_{I=1}^{N-1} \boldsymbol{W2_I}$ in the denominator is over all N individuals except for the individual carrying the private heterozygote A2/A3 itself.

This simple scoring scheme gives twice as much weight to the individuals homozygous for the allele A2 than to the individuals heterozygous for A2. Individuals that do not carry A2 are assigned score 0.

The probability $\boldsymbol{P3(L)}$ for a given individual L to have contributed the A3 allele to the private heterozygous genotype A2/A3 could be computed analogously. Thus, the probability that the private heterozygote A2/A3 at the site S represented by all three heterozygotes is a result of contamination involving individuals K and L (neither of them carrying A2/A3 genotype) could be computed as:

$$\boldsymbol{P23(K,L)} = \boldsymbol{P2(K)} * \boldsymbol{P3(L)} + \boldsymbol{P2(L)} * \boldsymbol{P3(K)}$$

where either $\boldsymbol{P2(K)} * \boldsymbol{P3(L)}$ or $\boldsymbol{P2(L)} * \boldsymbol{P3(K)}$ must be equal to 0 (both these terms could be positive only if the genotypes of both individuals K and L are A2/A3). The sum of $\boldsymbol{P23(K,L)}$ values over all pairs of individuals is equal to 1.

We computed the overall score (***H***) for the pair of individuals K and L by taking the sum of $\boldsymbol{P23(K,L)}$ values over all sites and dividing it by their number. This procedure was carried out for all possible pairs of individuals.

The sum of ***H***-scores over all pairs of the individuals is again equal to 1. Thus, the score ***H*** for a pair of individuals shows how likely it is that a hypothetical contamination involving this pair contributed to the pool of the sites with three heterozygous genotypes. ***H***-scores computed for all possible pairs of the individuals L4-L11 using custom Perl script are given in Supplementary Table 11. There are 28 pairwise combinations of the 8 individuals, which gives an expected value of score ***H*** equal to 1/28 = 0.0357.

If the sites carrying all three heterozygous genotypes were present among the sequenced individuals as a result of cross-sample contamination, we would expect to observe a limited number of pairs of individuals with extremely high ***H***-scores. However, the obtained ***H***-scores for all the pairs of the individuals were very similar (ranging from 0.0293 to 0.0413; Supplementary Table 11; Fig. 3c).

Apart from being hardly compatible with contamination, this result confirms that recombination signal in the data is not driven by any particular pair of individuals pointing to panmictic nature of the *A. vaga* population.

### XIV. Estimation of the transformation rate in the *A. vaga* population

Because organization of the *A. vaga* genome is incompatible with conventional meiosis, it is possible that genetic exchanges in the *A. vaga* population proceed through bacteria-like transformation involving homologous recombination. Under this scenario, population-scaled recombination rate $\mathbf{4N_e\,c}$ (where $\mathbf{N_e}$ is the effective population size and $\mathbf{c}$ is the probability of recombination between adjacent sites) would depend on the transformation frequency.

Estimates of population-scaled recombination rate ($\mathbf{4N_e\,c}$) could be inferred from the population genetics data and in turn could be used to obtain estimates of transformation rates. Therefore, we first derived an expression for $\mathbf{c}$, assuming that genetic exchange in *A. vaga* occurs by transformation.

Let us denote by $\mathbf{p(x)}$ the distribution of lengths of DNA segments which are transferred between genomes in the course of transformation. Then, the probability that any given genomic site would experience transformation in a single generation is given by:

$$\mathbf{a * \int_1^{\infty} x * p(x)dx} \quad (1)$$

where $\mathbf{a}$ is the per base probability of a transformation event, defined as **T/G**, where **T** is the expected number of transformation events per genome and **G** is the genome size in nucleotides.

Next, let us consider a pair of sites A and B residing $\mathbf{k}$ nucleotides apart from each other. Since a transfer of a DNA segment simultaneously spanning sites A and B would not lead to recombination between them, a recombination event between sites A and B would require a transformation event affecting only one of these two sites.

If sites A and B are separated by more than $\mathbf{x}$ nucleotides ($\mathbf{x < k}$), there could be no segments of the length $\mathbf{x}$ simultaneously spanning both sites. Consequently, in this case transformation with a segment of the length $\mathbf{x}$ affecting site A would always result in its recombination with site B. However, if the distance between A and B is less or equal to $\mathbf{x}$ ($\mathbf{x \geq k}$), then the probability that a segment of the length $\mathbf{x}$ does not cover B given that it covers A (and, thus, that recombination takes place) is $\mathbf{\frac{k}{x}}$.

Therefore, the probability of recombination between two sites given that one of the sites underwent transformation with the segment of the size $\mathbf{x}$ is:

$$\begin{cases} \mathbf{1, if\ x < k} \\ \mathbf{\frac{k}{x}, if\ x \geq k} \end{cases} \quad (2)$$

From (1) and (2) we can calculate the per generation probability of recombination between a pair of sites $\mathbf{k}$ nucleotides apart, $\mathbf{R(k)}$.

$$\mathbf{R(k) = 2a * \int_1^k x * p(x)dx + 2a * k * \int_k^{\infty} p(x)dx} \quad (3)$$

Equation (3) could be used to obtain the probability of recombination between two adjacent sites $\mathbf{c = R(1)}$:

$$\mathbf{c = R(1) = 2a} \quad (4)$$

In other words, with transformation the probability of recombination between adjacent sites is simply twice the per nucleotide per generation rate of transformation. Hence, the rate of recombination between adjacent sites can be used to infer the per generation rate of transformation.

The estimates of population-scaled recombination rate ($\boldsymbol{4N_e\,c}$) in the *A. vaga* population inferred from LD ($r^2$) decay and from the variance of pairwise differences were on the order of $10^{-2}$ (ranging from 0.015 to 0.046; see section «Estimation of the population-scaled recombination rate»). Assuming the effective size of the *A. vaga* population ($\boldsymbol{N_e}$ ) equal to $10^6$, this corresponds to **c**, as well as to $\boldsymbol{a}$ (4), of the order of $10^{-8}$. Given that the number of nucleotides in the haploid *A.vaga* genome is ~$10^8$, this translates to ~1 transformation event per generation.

Assuming that transformation is a means of interindividual genetic exchanges in *A. vaga*, the probability of recombination between a pair of sites is expected to increase with increasing distance between the sites, $\boldsymbol{k}$, only while $\boldsymbol{k}$ stays below the maximal length of the transferred segment, $\boldsymbol{L_M}$. This is due to the fact that for sufficiently large values of $\boldsymbol{k}$ ($\boldsymbol{k} > \boldsymbol{L_M}$), the probability of recombination between the pair of sites is the same, irrespective of the exact value of $\boldsymbol{k}$, as there are no more segments that can simultaneously span both sites. Patterns of LD decay observed in the data with LOESS estimates of $r^2$ reaching the values close to intercontig level already at 1,500 - 2,000 nucleotides (Fig. 2a) suggest that the characteristic size of the transferred segments also should be at least 1,500-2,000 nucleotides.

### XV. Estimation of the population-scaled recombination rate

To infer the population-scaled recombination rate ($\boldsymbol{4N_e\,c}$) in *A.vaga* from the rate of LD decay, we estimated the rate of short-range decay of $r^2$ with physical distance by applying a nonlinear regression model. This model is based on the equation for the expected value of $r^2$ under recombination-drift equilibrium with an adjustment for low mutation rate and sample size ($\boldsymbol{n}$) (ref. 28):

$$\boldsymbol{E(r^2) = \left[\frac{(10+C)}{(2+C)(11+C)}\right] * \left[1 + \frac{(3+C)(12+12C+C^2)}{n(2+C)(11+C)}\right]}.$$

where $\boldsymbol{C = 4N_e\,c_{sites}}$, $\boldsymbol{N_e}$ is the effective population size and $\boldsymbol{c_{sites}}$ is the recombination fraction between sites respectively.

To estimate the rate of LD decay, we used pairs of SNPs residing within the maximal distance of 500 bp from each other, with at least 4 copies of the minor allele among the individuals L4-L11. We fit nonlinear regression in R using the modified script by Marroni et al[19]. The estimated values of $\boldsymbol{4N_e\,c}$ based on the SNPs from the phased dataset 1 and based on the SNPs belonging to the more rigorously filtered phased dataset 2 were 0.0162 and 0.0150 respectively.

We also used the simpler formula for the expected value of $r^2$ without adjusting for mutations and sample size[20]. Under this basic model of recombination-drift equilibrium the expectation of $r^2$ is given by $\boldsymbol{E(r^2) = \left[\frac{1}{(1+C)}\right]}$. The estimates of $\boldsymbol{4N_e\,c}$ produced with the nonlinear regression based on this formula were 0.0265 and 0.0267 for the phased dataset 1 and 2 respectively.

Additionally, we estimated population-scaled recombination rate with Wakeley's moment method[21], as implemented in LDhat[16]. For this purpose, we first obtained Wakeley's estimates of recombination rate for all segments of the *A. vaga* genome harboring at least 5 phased non-singleton variants segregating among the individuals L4-L11 (n = 1,978). The Wakeley's estimate of population scaled recombination rate (expressed as $\boldsymbol{4N_e\,c}$), was obtained separately for each genomic segment and normalized by the segment size. Those regions that were identified as outliers for the normalized Wakeley's estimate of recombination rate using the interquartile range method were excluded from the further analyses (n = 254).

We further removed from consideration regions with less than 20 phased non-singleton variants, which left us with 258 genomic segments. The median length of the segments belonging to the resulting dataset is 872 base pairs and the median number of non-singleton phased variants harbored by these segments is 26.5. The median Wakeley's estimate of recombination rate across these 258 regions is 0.046, which is largely consistent with the estimates based on the rate of LD decay.

### XVI. Characterizing recombination in *A. vaga*

First, we investigated whether the probability of a recombination event depends on the GC-content of a genomic region.

As a proxy for recombination rate of a region we used three different measures:

1. Normalized minimum number of recombination events ($R_{min}$)
2. Normalized Wakeley's estimate of recombination rate
3. Fraction of the SNP pairs passing the modified four-gamete test

Estimation of the minimum number of recombination events ($R_{min}$) was performed for the same set of phased genomic segments (n = 258) which was utilized to obtain Wakeley's estimates (see section «Estimation of the population-scaled recombination rate»). Estimates of the minimum number of recombination events according to Hudson and Kaplan[13] for individual phased segments were inferred with LDhat[16]. We normalized the obtained values dividing them by the total number of non-singleton SNP pairs residing within the phased segment. Then, we computed the Pearson's correlation coefficient between the normalized minimum number of recombination events and GC-content of the genomic segment and observed that these two features are in a weak negative relationship (Pearson's correlation coefficient = -0.2; Extended Data Fig. 5a). Simple linear regression analysis showed that negative association between the probability of a recombination event and the GC-content of the region is significant (linear regression R-squared = 0.03998, regression coefficient = -0.056, P value for the slope = 0.0012). To confirm that the observed negative correlation is not an artifact of the larger sizes of the GC-poor segments, we also performed multiple linear regression analysis with GC-content and the size of the segment as the explanatory variables and normalized minimum number of recombination events as the response variable. The P value for the partial regression coefficient associated with the GC-content remained significant (partial regression coefficient = -0.058, P value = 0.000785).

Regions of low GC-content also showed a higher probability to experience a recombination event based on the estimates of recombination rate inferred with Wakeley's moment method (Pearson's coefficient of correlation between the population scaled normalized recombination rate and GC-content of the region = -0.13; linear regression R-squared = 0.0171, linear regression coefficient = -0.2593, P value for the slope = 0.0357).

In line with these observations, fractions of SNP pairs passing the modified four-gamete test confirmed the same pattern demonstrating a negative correlation with the GC-content of a contig (Extended Data Fig. 4a). Unlike analyses based on the minimum number of recombination events and Wakeley's estimate of recombination rate, which utilized individual phased genomic segments, this analysis was done in a contig-wise manner. For this purpose, we first retained only those contigs from the haploid sub-assembly that harbor at least 20 pairs of heterozygous SNPs (minor allele count ≥ 4) meeting conditions of the modified four-gamete test, which resulted in a set of 666 haploid contigs. Next, we computed fractions of SNP pairs passing the modified four-gamete test for each contig from the resulting set. Fraction of recombinant SNP pairs for these haploid contigs was negatively correlated with contig GC-content (Pearson's coefficient of correlation = -0.158; linear regression R-squared = 0.02496, linear regression coefficient = -1.7806, P value for the slope = 4.22e-05).

To confirm that the observed negative relationship is not due to larger sizes of GC-poor contigs, we uncoupled the effects of GC-content and contig size by comparing fractions of SNP pairs passing the modified four-gamete test among SNP pairs falling within the same distance bin for contigs of different GC-content (Extended Data Fig. 4b; Extended Data Fig. 5b). For this purpose, we subdivided contigs carrying heterozygous pairs of SNPs meeting the conditions of the modified four-gamete test (n=1,740) into 3 bins of approximately equal size according to their GC-content. Next, each considered pair of heterozygous SNPs was assigned to a group based on its distance bin and GC-content bin of the corresponding contig; proportions of recombinant SNP pairs were compared between 12 resulting groups (Extended Data Fig. 4b; Extended Data Fig. 5b).

Moreover, even if the contig identity was controlled for, recombination events showed tendency to occur in the GC-depleted regions of the contig. This has been demonstrated in the following way. For each contig of the haploid sub-assembly carrying pairs of SNPs passing the modified four-gamete test (MAC ≥ 4 among L4-L11), we selected the pair of SNPs separated by the smallest distance among all recombinant SNP pairs and located at least at a distance of 100 bp of each other (if there were SNP pairs satisfying these conditions). We treated intervals separating such sites (n = 1,014) as a proxy for the locations of recombination events (further referred to as recombinant intervals). To see if GC-content of such intervals is lower than would be expected by chance, we randomly sampled genomic intervals preserving the number, contig identity and the distribution of sizes of the actual recombinant intervals. This sampling procedure was repeated 1,000 times and the average GC-content of the recombinant intervals was compared to average values of GC-content for 1,000 random samples.

Reduction in GC-content of the recombinant intervals relative to random expectation was found to be significant (P = 0.019). A one-sided P value was computed as the fraction of 1,000 random interval datasets with an average GC-content lower or equal to the average GC-content of the recombinant intervals (Extended Data Fig. 4c).

To test if recombination is more likely to occur in genomic regions associated with particular annotation feature types, we compared proportions of recombinant SNP pairs among closely located SNP pairs falling within compartments of the genome annotated as exons or introns. To focus on the pairs of sites which could be unambiguously assigned to individual gene features, for this analysis we retained only closely located SNP pairs separated by no more than 200 base pairs and located within the same genomic compartment (exon or intron). To account for the difference in distribution of intra-pair distances between exonic and intronic SNPs, out of 10,817 pairs of heterozygous SNPs assigned to exons we selected 2,375 pairs matched for intra-pair distances to the 2,375 pairs of intronic SNPs. For this, we used the *matchit* function from the package *MatchIt* in R version 3.5.1. We applied the modified four-gamete test to these matched sets of exonic and intronic SNP pairs. This procedure allowed us to compare proportions of recombinant SNP pairs for exons and introns. Significance of the difference between proportions of recombinant SNP pairs among SNP pairs covered by exons and introns was assessed using two-sample Z-test for proportions as implemented in the *prop.test* function in R employed without continuity correction (Extended Data Fig. 4d).

### XVII. Analysis of correlations between distances separating haplotypic counterparts in pairs of individuals

To analyze how distances between haplotypic counterparts are correlated for different pairs of individuals L4-L11, we focused on the same set of phased genomic segments that was used in previous analyses (n=262, see section "Signatures of recombination within individual phased genomic regions" in Methods and Supplementary Methods IX) additionally requiring the minimum span of 500 base pairs. This resulted in the set of 244 segments carrying at least 20 non-singleton SNPs simultaneously phased in all individuals L4-L11. For each such segment, we reconstructed sequences of the two haplotypes in each individual based on the corresponding sequence from the haploid sub-assembly and the set of phased non-singleton SNPs using BCFtools v.1.4.1. Only non-singleton SNPs that satisfied all filtering criteria and were simultaneously phased in L4-L11 were used to reconstruct haplotypes; the remaining sites were treated as monomorphic.

Next, for each pair of individuals (further on referred to as "individual 1" and "individual 2"), we retained only those segments where it was possible to assign 4 haplotypes into 2 pairs, such that haplotypes within the same pair likely had a recent common ancestor. For this purpose, for each segment, we first computed a proportion of nucleotide differences for all pairwise comparisons between reconstructed sequences of haplotypes. Next, for each haplotype in individual 1, we identified the more and the less similar haplotype in individual 2. Cases where it was not possible to unambiguously identify the 'best hit' of each haplotype were discarded. For this, we compared the minimal ($N_{MIN}$) and the maximal ($N_{MAX}$) number of nucleotide differences separating the considered haplotype of individual 1 from the two haplotypes in individual 2. The haplotype was defined as having an unambiguous 'best hit' in individual two if the difference between $N_{MAX}$ and $N_{MIN}$ was 2 SNPs or more ($N_{MAX}$-$N_{MIN} \geq 2$). Analogously, for each haplotype in individual 2, we identified the 'best hit' in individual 1.

Finally, we retained only those segments where haplotypes could be assigned into reciprocally best matching pairs. This procedure aims at minimizing the effects of gene conversion, as an event of gene conversion private to one of the individuals makes both haplotypes of this individual similar to the same haplotype of individual 2. In this case, haplotypes can no longer be assigned into reciprocally best matching pairs and such segments are discarded from analysis.

As a result of the above-described procedure, for each pair of individuals, we identified a subset of phased segments where 4 haplotypes from two individuals were assigned into unambiguous pairs of haplotypic counterparts. Then, for each such segment, we selected the minimal and the maximal distance separating haplotypes within the same pair and examined whether these distances were correlated. We found that for all pairs of individuals the distances separating haplotypic counterparts were significantly correlated (Supplementary Table 13).

It is conceivable that shared events of gene conversion could result in correlation between the distances separating haplotypic counterparts. To address this possibility, we examined the relationship between the minimal/maximal distance in pairs of haplotypic counterparts and distances between the two haplotypes within the same individual. If correlations between minimal and maximal distances separating haplotypic counterparts were due to gene conversion, we would expect both these distances to be correlated with within-individual interhaplotypic distances.

Although within-individual interhaplotypic distances were usually positively correlated with the maximal distance separating haplotypic counterparts from different individuals (Supplementary Table 14), in most cases there was no significant correlation between within-individual haplotypic distances and the minimal distance separating haplotypic counterparts (Supplementary Table 14). We further confirmed that the correlation between distances separating haplotypic counterparts is not explained by gene conversion by applying multiple linear regression. For this purpose, for each pair of individuals, the maximal distance separating haplotypic counterparts from different individuals was treated as the response variable, and the minimal distance separating haplotypic counterparts, as well as within-individual distances between haplotypes were treated as the explanatory variables. The P values on partial t-tests on minimal distance separating haplotypic counterparts remained significant (Supplementary Table 15), confirming that joint inheritance of both haplotypes is the likely explanation of the observed correlation between interhaplotypic distances in pairs of individuals.

### Supplementary Note

Collectively, the presented data provide solid evidence for recombination in the population of *A. vaga*. Still, the signal of recombination could be driven by gene conversion. Because some of the techniques commonly used to infer recombination are sensitive to gene conversion, we devised tests to distinguish signatures of recombination involving genetic exchanges from those of gene conversion.

Decay of LD measured in a conventional way (Fig. 2a, b; Extended Data Fig. 2a, b, c), as well as recombination signal inferred from the sum of the distances and PHI tests (Fig. 3d), conceivably could be associated with gene conversion. To address this, we devised the modified four-gamete test (Fig. 2e; Supplementary Methods VIII) and showed that the observed decline in LD with physical distance could not be explained by gene conversion alone (Fig. 2f, g).

Incongruent phylogenies of different genomic regions observed in consensus networks built from the unphased genotype data (Fig. 3e; Extended Data Fig. 3c, d; Supplementary Methods X) could also possibly be caused by gene conversion. However, phylogenetic analysis of individual haplotypes at different loci revealed patterns that could only result from genetic exchanges (Extended Data Table 1; Fig. 4a, b, c, d; Methods). Namely, we observed multiple cases when the two haplotypes of a single individual had closest haplotypic counterparts in two different individuals. Gene conversion can introduce incongruence to phylogenies of haplotypes by increasing the similarity of the two haplotypes of a single individual to each other. However, it cannot increase the similarity between haplotypes harbored by different individuals[22].

Still, gene conversion can create spurious clustering of haplotypes from a single individual with the two haplotypes from different individuals[23]. To see this, consider two closely related individuals A and B (carrying haplotypes hapA.1/hapA.2 and hapB.1/hapB.2 respectively; Supplementary Figure 1), and the third individual C (with haplotypes hapC.1/ hapC.2) which is more distantly related to A and B. Under obligate asexual reproduction in the absence of gene conversion, we would expect the haplotypic counterparts from individuals A and B to be clustered together, forming two pairs of similar haplotypes[24] hapA.1-hapB.1 and hapA.2-hapB.2 (the existence of such clustering in asexuals has been suggested by M. Meselson and is commonly referred to as the 'Meselson effect'[24]; Supplementary Figure 1). However, if gene conversion replaced the sequence of one of the haplotypes in the individual B with the sequence of the other (converting hapB.1 to hapB.2), there would no longer exist a haplotypic counterpart of hapA.1 in individual B (Supplementary Figure 1). In this case, one haplotype of individual A would be most similar to a haplotype from B (hapA.2-hapB.2), while the other, to a haplotype from individual C (hapA.1-hapC.1), as the corresponding haplotype in B had been replaced by the sequence of its allelic counterpart.

However, the resulting phylogenetic signal of gene conversion can be distinguished from the signal of genetic exchange. In contrast to genetic exchange, gene conversion would make the distance separating the two haplotypes of individual B (hapB.1-hapB.2) shorter than the distances separating each of these haplotypes from its closest haplotypic counterpart in another individual (Supplementary Figure 1). The cases in which this condition does not hold cannot arise from gene conversion, and by exclusion, have to be due to genetic exchange.

We employed this logic to identify cases of putative genetic exchange (see Methods; Extended Data Table 1; Fig. 4a, b, c, d) in the phased haplotype data and found a signature of genetic exchange in 56 out of the 262 phased segments used for the analysis (only cases with bootstrap support ≥ 70% were considered, see Methods).

The analyzed phased segments cover a small portion of the *A. vaga* genome (260,204 bp out of 76,679,421 bp included in the haploid sub-assembly). This is due to a high cutoff set on the number of SNPs simultaneously phased in all individuals L4-L11 (we required a minimum of 20 such SNPs for a segment to be included in this analysis; see Methods). However, it is likely that a substantial fraction of the *A. vaga* genome has been shaped by genetic exchange, as ~21% of the analyzed segments show a signature of this process.

Finally, we corroborated all previous findings through analysis of triallelic sites. We found a substantial number of triallelic sites harboring all three possible heterozygous genotypes. This observation cannot be attributed to action of gene conversion, as there is no possible way in which three heterozygotes can emerge from this process. Nevertheless, sites carrying all three heterozygous genotypes can arise through recurrent or back mutations without genetic exchanges. However, we showed that the observed number of such sites is significantly larger than that expected due to recurrent or back mutations (see Main Text; Fig. 3b, c; Supplementary Table 9; Supplementary Methods XII).

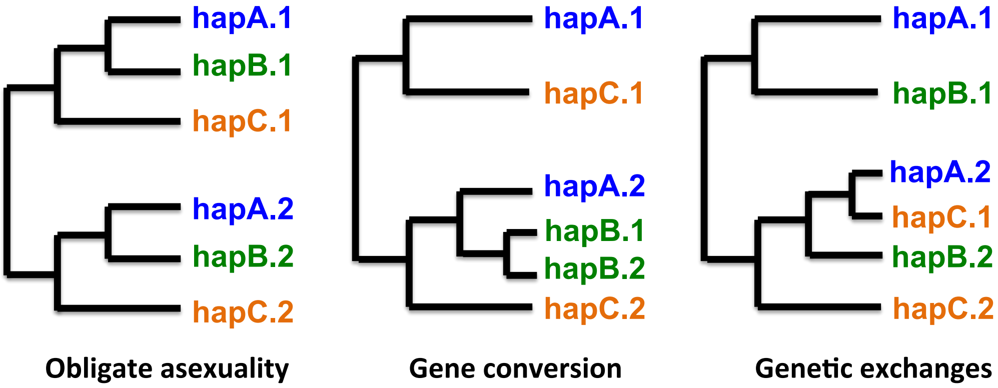

**Supplementary Figure 1 Patterns of relationship between haplotypes from different individuals expected under different evolutionary scenarios[22,24].** Indices 1 and 2 designate the two haplotypes of a single individual.

Thus, the presented data are not compatible with strictly asexual reproduction in *A. vaga,* with or without gene conversion. Although the apparent lack of males and homologous chromosomes[1] virtually ruled out conventional meiosis in bdelloid rotifers, it still remained unclear whether they engage in genetic exchanges through an atypical version of meiosis observed in a few plants (*Oenothera*-like meiosis)[22,23] as proposed by Signorovitch *et al.* (2015, 2016), or through transformation[25] as proposed by Debortoli *et al.* (2016). Many of the patterns observed in our study and suggestive of genetic exchange can emerge under both these scenarios:

1) Obviously, both *Oenothera*-like meiosis and transformation can lead to coexistence of all three heterozygous genotypes at triallelic sites.

2) Unlike transformation, *Oenothera*-like meiosis alone is not expected to cause a decline in LD with distance as it involves no crossing over. However, as discussed in the main text, gene conversion between allelic regions would suffice to explain a decay of LD irrespective of the mode of reproduction.

3) Pairs of sites passing the modified four-gamete test in principle could also arise under *Oenothera*-like meiosis as well as under transformation. Obviously, transfer of DNA segments occurring during transformation can produce a pair of individuals, each heterozygous at two loci, carrying all four haplotypes (Fig. 2e; Supplementary Figure 2a). On the contrary, gene conversion alone cannot give rise to such a pair. This is the basis of the modified four-gamete test employed in this study to distinguish gene conversion from genetic exchange (Supplementary Methods VIII). Nevertheless, homologous recombination during genetic exchange is not a prerequisite for the existence of pairs of sites passing the modified four-gamete test. Indeed, gene conversion in conjunction with genetic exchange in the form of *Oenothera*-like meiosis would suffice to explain the existence of such pairs of sites (Supplementary Figure 2b).

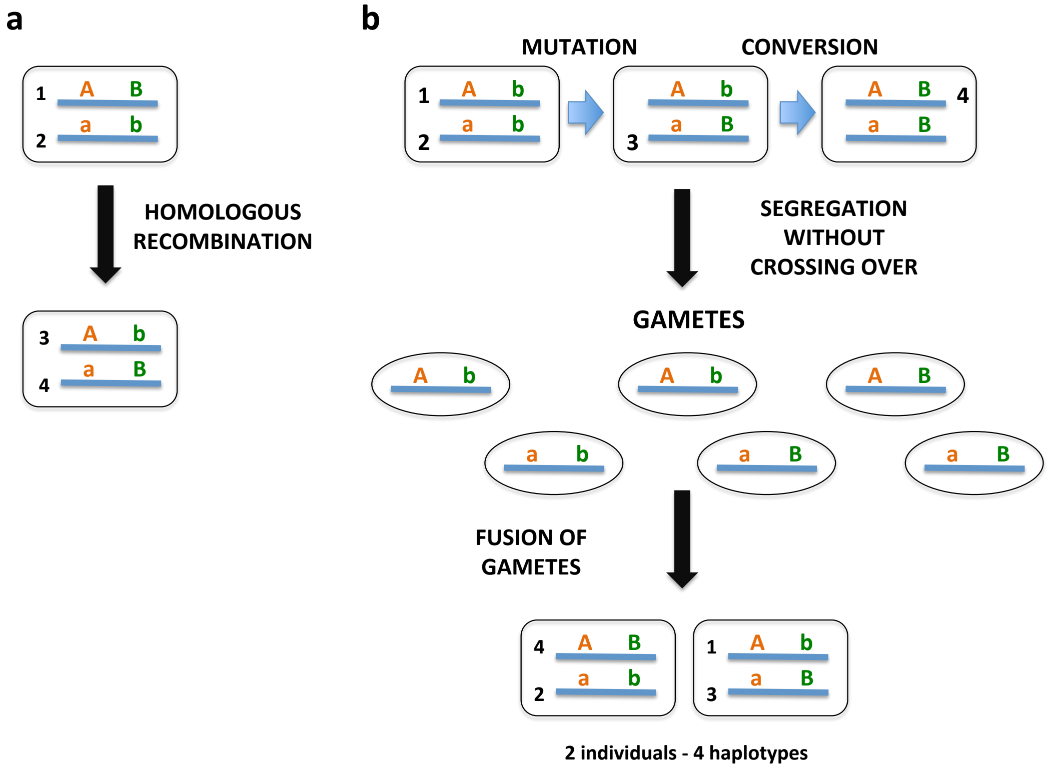

**Supplementary Figure 2 Emergence of four haplotypes for a pair of heterozygous sites in two individuals due to homologous recombination during transformation (a) or due to gene conversion in conjunction with genetic exchanges in the form of *Oenothera*-like meiosis (b).**

To distinguish between these two modes of genetic exchange in *A. vaga,* we employed phylogenetic analysis of haplotypes. Although both *Oenothera*-like meiosis and transformation will lead to incongruence between phylogenies of the two haplotypes[22], these two scenarios could be discriminated by comparing patterns of incongruence at different loci. Under *Oenothera*-like meiosis, all chromosomes forming the same complex[22] are passed on together from generation to generation. In this case, segregation with subsequent fusion of gametes will lead to different evolutionary histories of the two haplotypes found within a single individual. However, as haplotypes belonging to the same complex are inherited together, the patterns of incongruence between the two haplotypes at different loci are expected to be identical[22]. In contrast, transfer of DNA segments is expected to affect remote genomic regions independently of one another, which should lead to different patterns of incongruence at different loci.

In our data, we found multiple cases when the two haplotypes of a single individual clustered with haplotypes from two different individuals (Extended Data Table 1; Fig. 4a, b, c, d). However, these pairs of individuals were usually different for different loci, so that each individual exhibited multiple patterns of incongruence between the two haplotypes (Extended Data Table 1; Fig. 4a, b, c, d).

Moreover, for 25 out of 28 possible pairs of individuals L4-L11, we found cases when there existed a third individual harbouring one haplotype clustered with a haplotype of the first individual from the pair and another haplotype clustered with a haplotype of the second individual (Supplementary Table 12; only cases with bootstrap support $\geq 70\%$ were considered, see Methods). In other words, many distinct patterns of incongruence were present in the data and these patterns are not likely to arise from gene conversion (see above).

Another line of evidence against *Oenothera*-like meiosis as the main mode of genetic exchange in *A. vaga* comes from the analysis of correlations between distances separating haplotypic counterparts for different pairs of individuals (Supplementary Methods XVII; Extended Data Fig. 6a, b, c; Supplementary Tables 13-15). The rationale behind this analysis is as follows. *Oenothera*-like meiosis leads to independent inheritance of the two complexes of haplotypes[22]. If this type of meiosis is frequent, evolutionary histories of two different complexes within the individual are usually different, and, consequently, distances between haplotypes belonging to two separate complexes for a pair of individuals are not expected to be correlated. However, we found that for all pairs of individuals, the distances separating haplotypic counterparts were significantly positively correlated (Supplementary Table 13; Extended Data Fig. 6a, b, c), and these correlations were not due to gene conversion (Supplementary Methods XVII; Supplementary Tables 14-15).

In summary, although our data do not exclude the possibility of rare occurrences of *Oenothera*-like meiosis in bdelloid rotifers, such meiosis alone is not sufficient to explain our data. By contrast, it is sufficient to assume the presence of a single mechanism of genetic exchange introducing different patterns of incongruence at different loci such as transformation to explain our findings.

### Supplementary Tables

**Excel file containing Supplementary Tables 1-2, 5-9 and 11-15 arranged as separate Excel worksheets can be accessed at:**
https://figshare.com/s/3f6d81a9cbce5e0b619e.

**Supplementary Table 1. Summary statistics on whole-genome sequencing of 11 wild-caught *A. vaga* individuals.** Coverage for each individual was determined based on alignments of the 2×100 bp HiSeq paired-end reads against the initial *A. vaga* genome assembly and against the haploid sub-assembly. Alignments were performed with Bowtie 2. Paired-end reads mapping to no more than two positions in the initial assembly were aligned against the haploid sub-assembly. The resulting alignments were additionally filtered (see Supplementary Methods III), and only reads uniquely mapped to the haploid sub-assembly were retained. Statistics of coverage for the haploid sub-assembly are based on the final filtered alignments.
*In the accompanying Excel file.*

**Supplementary Table 2. Assembly statistics for the obtained *A. vaga* reference genome (L1).** The statistics were generated with QUAST v5.0.0 and, unless noted otherwise, are based on contigs (or haploid segments in case of haploid sub-assembly) with a minimum length of 500 bp.
*In the accompanying Excel file.*

**Supplementary Table 3. Estimates of genomic divergence of the sequenced *A. vaga* individuals from the published *A. vaga* genome.** For each sequenced individual, 1,000,000 reads were randomly drawn and used for the BLAST search against the published *A. vaga* genome assembly. For all reads with reported hits in the *A. vaga* genome assembly, the best *blastn* hit was chosen, and average (or median) nucleotide identity was computed across these hits.

| Sample | Average identity, % | Median identity, % |
|---|---|---|
| **L1** | 87.35 | 87.76 |
| **L2** | 87.38 | 87.76 |
| **L3** | 87.52 | 87.76 |
| **L4** | 87.43 | 87.76 |
| **L5** | 87.5 | 87.76 |
| **L6** | 87.47 | 87.76 |
| **L7** | 87.49 | 87.76 |
| **L8** | 87.4 | 87.76 |
| **L9** | 87.17 | 87.36 |
| **L10** | 87.45 | 87.76 |
| **L11** | 87.33 | 87.64 |

**Supplementary Table 4. Numbers of genomic sites with called genotypes in the raw SNP datasets, and numbers of sites retained after applying various quality filtering steps.** Numbers of sites included in the final filtered datasets (stringent SNP datasets I and II) and used throughout the analyses are shown in bold.

| | SNP dataset | |
|---|---|---|
| | **Dataset I (variable sites only)** | **Dataset II (variable and invariant sites)** |
| **Prior to filtration** | 3,318,352 | 76,306,143 |
| **Filter:** | | |
| **Sites within 10 bp of an indel removed** | 2,979,193 | 76,004,960 |
| **Sites with missing genotypes and sites with QUAL <50 removed** | 2,655,917 | 49,222,726 |
| **Sites on haploid segments shorter than 1,000 bp removed** | 2,634,341 | 48,730,324 |
| **Sites residing within repetitive regions removed** | 2,596,490 | 47,531,499 |
| **Sites covered by <10 reads in any of the samples removed** | 2,409,323 | 44,541,675 |
| **Sites with extremely high or low coverage removed** | 2,391,710 | 43,924,505 |
| **Sites within the windows outliers for SNP density removed** | **2,282,099** | **42,850,155** |

**Supplementary Table 5. Pairwise genotypic distances between the sequenced *A. vaga* individuals.** Genotypic distances were computed based on the sites of the haploid sub-assembly simultaneously called in all sequenced individuals L1-L11. Only monomorphic and biallelic sites from the stringent SNP dataset II were used in the analysis. For a pair of *A. vaga* individuals, genotypic distance was calculated for each site as the difference in the number of non-reference variants (0, 1 or 2), and then averaged over all analyzed sites. Distances between the individuals belonging to different clusters are highlighted in blue, and distances within the small and the large cluster are shown in red and green respectively.
*In the accompanying Excel file.*

**Supplementary Table 6. Statistics on the span of phased blocks assembled with HapCUT2 and included in the phased dataset 1.** Homozygous SNPs were appended to phased blocks based on the block assignment of closest flanking heterozygous SNPs.
*In the accompanying Excel file.*

**Supplementary Table 7. Whole-genome numbers of phased SNPs belonging to phased blocks assembled with HapCUT2 and included in the phased dataset 1.** Total numbers of phased SNPs were determined as the numbers of SNPs belonging to any phased block encompassing at least 2 heterozygous SNPs irrespective of its length. Homozygous SNPs were appended to phased blocks based on the block assignment of closest flanking heterozygous SNPs.
*In the accompanying Excel file.*

**Supplementary Table 8. Testing for Hardy-Weinberg equilibrium.** Exact tests of HWE were performed on biallelic SNPs from the stringent SNP dataset II simultaneously called in all sequenced individuals L1-L11. Tests were performed separately within the large cluster, L4-L11, and together for all sequenced individuals, L1-L11. Analysis was based on the sets of SNPs common (MAC $\geq$ 4) among L4-L11 or L1-L11. SNPs with the exact test P value $\leq$ 0.05 were considered to be out of HWE. Statistics on the observed to expected ratios of genotype counts were obtained for the sites with unambiguously identified major and minor alleles.
*In the accompanying Excel file.*

**Supplementary Table 9. Observed and expected numbers of triallelic sites harboring all three possible heterozygous genotypes among the sequenced individuals.** For each dataset, the difference between observed and expected numbers of triallelic sites harboring all three possible heterozygous genotypes was significant at P < 2.2e-16 (one sample Z-test for proportions, highlighted in yellow).
*In the accompanying Excel file.*

**Supplementary Table 10. Per individual numbers of sites with a unique heterozygous genotype private to the given individual among triallelic sites carrying all three heterozygous genotypes.** Only those triallelic sites harboring all three heterozygous genotypes among L4-L11 for which exactly one private heterozygous genotype exists (n = 607) were considered.

| Sample | Number of sites with a unique heterozygous genotype private to the given individual among the sites harboring three heterozygotes |
|---|---|
| L4 | 77 |
| L5 | 77 |
| L6 | 88 |
| L7 | 76 |
| L8 | 79 |
| L9 | 74 |
| L10 | 72 |
| L11 | 64 |

**Supplementary Table 11. Pairwise scores (*H*-scores) reflecting the relative probability to observe a triallelic site harboring all three heterozygotes as a result of hypothetical exchange or contamination involving this pair of individuals.** Only those triallelic sites harboring all three heterozygous genotypes among L4-L11 for which a single private heterozygous genotype exists (n = 607) were considered.
*In the accompanying Excel file.*

**Supplementary Table 12. Patterns of incongruence observed for different phased segments of the *A. vaga* genome.** For each pair of individuals L4-L11, we computed the number of phased segments (out of 262 analyzed segments) when there existed a third individual most similar to the first individual from the pair with respect to one haplotype and most similar to the second individual from the pair with respect to the other haplotype (only cases with bootstrap support values ≥ 70% were considered, see Methods and Extended Data Table 1). Some segments exhibited the above-described pattern for more than one pair of individuals.
*In the accompanying Excel file.*

**Supplementary Table 13. Correlations between minimal and maximal distance separating haplotypic counterparts in different pairs of individuals L4-L11.** Distances expressed as proportions of nucleotide differences were computed for the segments of the *A. vaga* genome simultaneously phased in all individuals L4-L11 and carrying at least 20 non-singleton SNPs. For each comparison of individuals, we utilized only those segments where 4 haplotypes from two individuals could be unambiguously assigned into 2 pairs, each pair comprising a haplotype from the first individual and its counterpart from the second individual. Haplotypic counterparts assigned to the same pair are likely to share a recent common ancestor. For each segment, we selected the minimal and the maximal distance separating haplotypic counterparts in two individuals (see Supplementary Methods XVII). For each comparison of individuals, Pearson's r between minimal and maximal distance separating haplotypic counterparts in different phased segments is shown. Corresponding P values are given in parentheses.
*In the accompanying Excel file.*

**Supplementary Table 14. Correlations between distances separating the two haplotypes within a single individual and interindividual haplotypic distances.** Distances expressed as proportions of nucleotide differences were computed for the segments of the *A. vaga* genome simultaneously phased in all individuals L4-L11 and carrying at least 20 non-singleton SNPs. For each comparison of individuals, we utilized only those segments where 4 haplotypes from two individuals could be unambiguously assigned into 2 pairs, each pair comprising a haplotype from the first individual and its counterpart from the second individual. Haplotypic counterparts assigned to the same pair are likely to share a recent common ancestor. For each segment, we selected the minimal and the maximal distance separating haplotypic counterparts in two individuals (see Supplementary Methods XVII). For each comparison of individuals, Pearson's r between distance separating the two haplotypes within the first individual in the pair and minimal/maximal distance to the haplotypic counterpart in the second individual is shown. Corresponding P values are given in parentheses. For each interindividual comparison, the first individual in the pair is the one listed in the first column.
Cases where both correlations were significant are shown in bold.
*In the accompanying Excel file.*

**Supplementary Table 15. Multiple linear regressions for maximal distances separating haplotypic counterparts in two individuals.**

For each pair of individuals, the maximal distance separating haplotypic counterparts from different individuals was treated as the response variable, while the minimal distance separating haplotypic counterparts, as well as distances between the two haplotypes of a single individual were treated as the explanatory variables (see Supplementary Methods XVII). Values of R-squared with P values from partial t-tests in parentheses are shown. P values on partial t-tests on different explanatory variables are given in the following order: 1) minimal distance between haplotypic counterparts, 2) distance between the two haplotypes within the first individual, 3) distance between the two haplotypes within the second individual.

For each interindividual comparison, the first individual in the pair is the one listed in the first column.

*In the accompanying Excel file.*
